# XIAP controls RIPK2 signaling by preventing its deposition in speck-like structures

**DOI:** 10.1101/545400

**Authors:** Kornelia Ellwanger, Christine Arnold, Selina Briese, Ioannis Kienes, Jens Pfannstiel, Thomas A. Kufer

## Abstract

The receptor interacting serine/threonine kinase 2 (RIPK2) is essential for linking activation of the pattern recognition receptors NOD1 and NOD2 to cellular signaling events.

Recently, it was shown that RIPK2 forms higher order molecular structures *in vitro*, which were proposed to activate signaling. Here, we demonstrate that RIPK2 forms detergent insoluble complexes in the cytosol of host cells upon infection with invasive enteropathogenic bacteria. Formation of these structures occurred after NF-κB activation and depends on the CARD of NOD1 or NOD2. Complex formation upon activation was dependent on RIPK2 autophosphorylation at Y474 and influenced by phosphorylation at S176. Inhibition of activity of the cIAP protein XIAP induced spontaneous complex formation of RIPK2 but blocked NOD1-dependet NF-κB activation. Using immunoprecipitation, we identified 14-3-3 proteins as novel binding partners of non-activated RIPK2, whereas complexed RIPK2 was bound by the prohibitin proteins Erlin-1 and Erlin-2.

Taken together, our work reveals novel roles of XIAP, 14-3-3 and Erlin proteins in the regulation of RIPK2 and expands our knowledge on the function of RIPK2 posttranslational modifications in NOD1/2 signaling.

## INTRODUCTION

The first line of defense in innate immunity in mammals encompasses several groups of pattern recognition receptors (PRRs) including Toll-like receptors (TLRs), cytosolic RIG-like helicases (RLHs), and NOD-like receptors (NLRs), recognizing pathogen-associated molecular pattern (PAMPs) (Dostert, Meylan et al., 2008, Janeway, 1989). Most human NLRs are involved in innate and adaptive immunity via transcriptional regulation of MHC class I and class II or regulating the innate immune response (Ting, Lovering et al., 2008). The NLR proteins NOD1 and NOD2 are intracellular patter-recognition receptors (PRR), sensing bacterial peptidoglycan (PGN)-derived y-D-glutamyl-meso-diaminopimelic acid (iE-DAP) and muramyl dipeptide (MurNAc-L-Ala-D-isoGln, MDP), respectively (Chamaillard, Hashimoto et al., 2003, Girardin, Boneca et al., 2003a, Girardin, Boneca et al., 2003b, Inohara, Ogura et al., 2003). Activation of both, NOD1 and NOD2 induces the NF-κB pathway (Girardin, Tournebize et al., 2001, Ogura, Inohara et al., 2001) by recruitment of the adaptor protein receptor interacting serine/threonine kinase 2 (RIPK2) (Girardin et al., 2001, Inohara, Koseki et al., 2000). RIPK2 (RIP2/RICK/CARDIAK) RIPK2 belongs to the RIPK family, a group of serine/threonine protein kinases (Navas, Baldwin et al., 1999, Thome, Hofmann et al., 1998). However, RIPK2 lacks the RHIM domain found in the cell death associated members RIPK1 and RIPK3 (Humphries, Yang et al., 2015). RIPK2 is essential for NOD1-and NOD2-mediated NF-κB activation and can contribute to T cell activation and apoptosis (Hall, Wilhelm et al., 2008, Nachbur, Stafford et al., 2015, Ruefli-Brasse, Lee et al., 2004, Tigno-Aranjuez, Benderitter et al., 2014).

The interaction of NOD1 and NOD2 with RIPK2 is mediated by heterotypic CARD:CARD interactions, involving residues in the exposed surfaces of the CARD domains of RIPK2 and NOD1/2 (Maharana, Pradhan et al., 2017, Manon, Favier et al., 2007, Mayle, Boyle et al., 2014). *In vitro*, this can result in stable rope-like structures, which were recently proposed to be platforms for subsequent NF-κB activation (Gong, Long et al., 2018, Pellegrini, Desfosses et al., 2018).

RIPK2 is controlled by complex posttranslational modification events, including autophosphorylation at several sites (Dorsch, Wang et al., 2006, Pellegrini, Signor et al., 2017, Tigno-Aranjuez, Asara et al., 2010). Best described are the phosphorylation events at S176 and Y474, which are associated with activity and structural changes (Pellegrini et al., 2017). The role and outcome of these phosphorylation events is not entirely understood. On the one hand side, it was shown that kinase activity of RIPK2 is dispensable for signaling and might only affect protein stability (Abbott, Wilkins et al., 2004, Windheim, Lang et al., 2007). On the other hand, additionally to resulting in protein instability, inhibition of kinase activity by the tyrosine kinase inhibitors Gefitinib and Erlotinib or the RIPK2 specific compounds WEHI-345 and GSK583 was shown to reduce signaling (Haile, Votta et al., 2016, Nachbur et al., 2015, Tigno-Aranjuez et al., 2010). Some insight on this controversy was provided by the recent identification that RIPK2 inhibitors can also block interaction of RIPK2 with the E3 ubiquitin ligase X-linked inhibitor of apoptosis (XIAP), which is essential for RIPK2-mediated NF-κB activation (Goncharov, Hedayati et al., 2018). RIPK2 is modified by K63-, K27 and M1-linked ubiquitination at K209, located in its kinase domain (Hasegawa, Fujimoto et al., 2008, Panda & Gekara, 2018). XIAP is the essential E3 for RIPK2 ubiquitination and interacts with RIPK2 through its baculoviral IAP-repeat (BIR) 2 domain (Krieg, Correa et al., 2009). XIAP also ubiquitinates K410 and K538 with K63-linked ubiquitin, which was shown to be important for NOD2 signaling (Goncharov et al., 2018). XIAP binding to RIPK2 recruits the linear chain assembly complex (LUBAC) (Damgaard, Nachbur et al., 2012). Moreover, further E3 ubiquitin ligases, including cellular inhibitor of apoptosis 1 (cIAP1) and cIAP2 (Bertrand, Doiron et al., 2009), TNF receptor-associated factor (TRAF) 2 and TRAF5 (Hasegawa et al., 2008), and ITCH (Tao, Scacheri et al., 2009) were shown to participate in RIPK2 ubiquitination. However, their physiological roles remain to be clarified. Ubiquitination of RIPK2 leads to recruitment of the transforming growth factor β-activated kinase 1 (TAK1). This ultimately triggers activation of the IκB kinase (IKK) complex (Hasegawa et al., 2008) and MAPK signaling (Girardin et al., 2001).

Here, we provide novel insights into RIPK2 biology. We found RIPK2 to form high molecular weight complexes, that we termed RIPosomes, in the cytosol of epithelial cells upon infection with invasive bacterial pathogens such as *Shigella flexneri* and enteropathogenic *Escherichia coli*. Upon bacterial infection, we observed localization of RIPK2 to the bacterial entry sites, followed by formation of dot-like structures and a changed electrophoretic mobility of RIPK2. Complex formation occurred after NF-κB activation, was dependent on the CARD of NOD1 or NOD2, and autophosphorylation of RIPK2 Y474. XIAP knockdown induced sterile RIPosome formation. Finally, we identified novel binding partners of the soluble and complexed forms of RIPK2 and showed that 14-3-3 proteins and Erlin proteins differentially bound these. Our data provide novel insights into the functional consequences of phosphorylation and ubiquitination events in RIPK2 and show that RIPosomes are not associated with immediate early signaling of NOD1.

## RESULTS

### RIPK2 forms cytosolic RIPosomes upon bacterial infection

To study details of RIPK2 function, we generated a stable inducible HeLa cell line expressing human EGFP-RIPK2. In order to activate RIPK2, we used the invasive gram-negative bacterium *S. flexneri* that physiologically activates the NOD1 pathway (Girardin et al., 2001). Infection of EGFP-RIPK2 cells with the invasive *S. flexneri* M90T strain increased IL-8 secretion compared to cells exposed to the non-invasive *S. flexneri* strain BS176 or EGFP expressing control cell line (Fig.1A). Albeit overexpression of RIPK2 led to some NF-κB activation and induction of IL-8 (McCarthy, Ni et al., 1998), the inflammatory response 6 h p.i. was much higher, also in cells induced for EGFP-RIPK2 expression for 16 h, compared to non-induced cells (Fig.1A).

**Figure 1:**
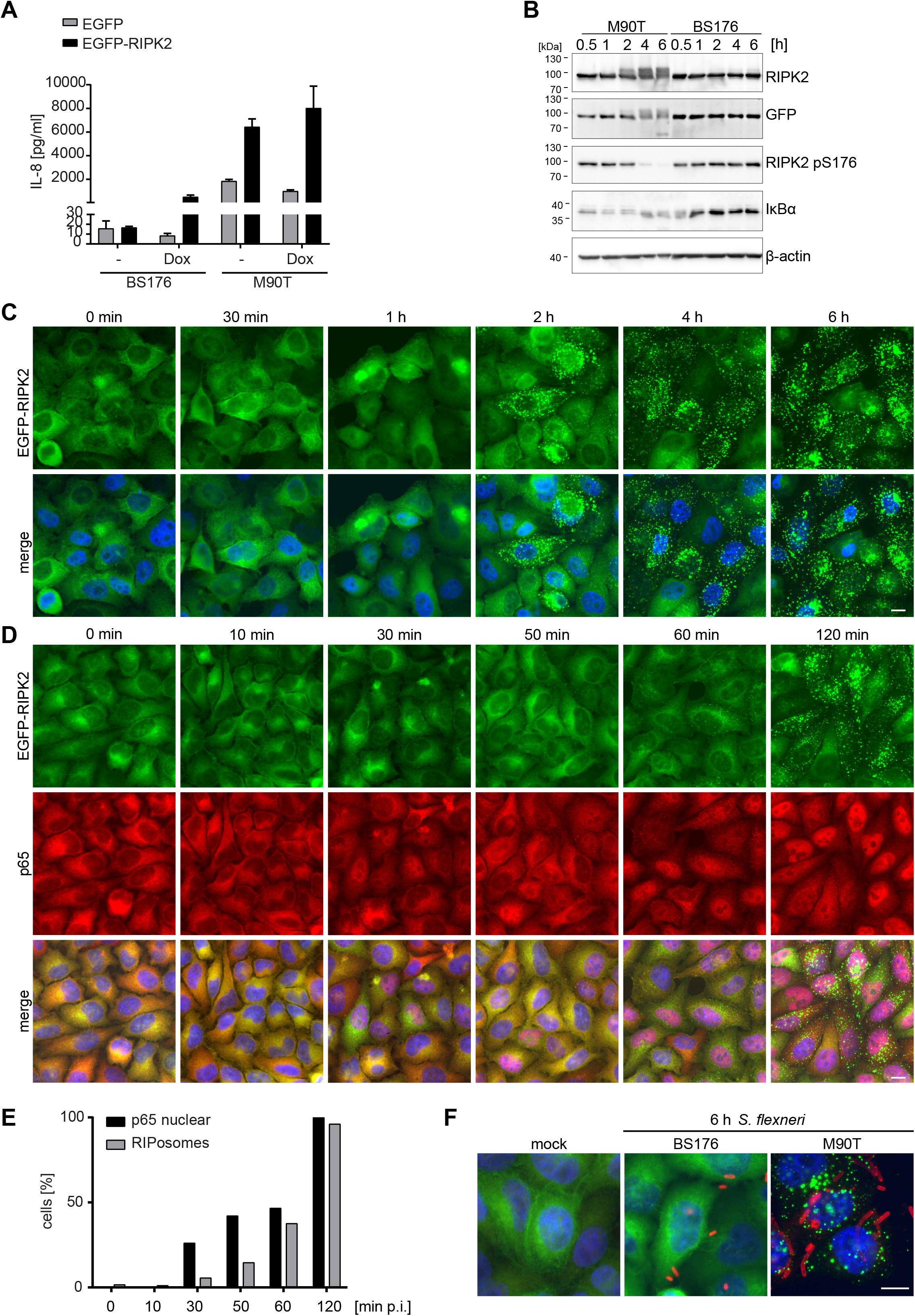
RIPK2 forms RIPosomes upon *Shigella* infection. **A:** IL-8 secretion of HeLa EGFP-RIPK2 and HeLa EGFP cells 6 h post infection with *S. flexneri* M90T or BS176. Mean + S.D. of one representative experiment conducted in triplicates is shown. **B:** Immunoblot analysis of *S. flexneri* M90T infected HeLa EGFP-RIPK2 cells infected with *S. flexneri* M90T for the indicated time. **C:** Indirect immunofluorescence micrographs of HeLa EGFP-RIPK2 cells infected with *S. flexneri* M90T for the indicated time. EGFP-RIPK2 signal and a merge with DNA staining is shown. Scale bar = 10 µm. **D:** Indirect immunofluorescence micrographs of HeLa EGFP-RIPK2 cells infected for the indicated time with *S. flexneri* M90T. Signals for p65 and EGFP-RIPK2 and a merge with DNA staining is shown. Scale bar = 10 µm. **E:** Left panel: Quantification of RIPosomes and nuclear p65 staining in the cells from (D). In total 200 cells were quantified. Right panel: Indirect immunofluorescence micrographs of HeLa EGFP-RIPK2 cells infected with *S. flexneri* M90T and BS176 for the indicated time. Signal of an anti-*Shigella*-LPS 5a antibody and EGFP-RIPK2 together with DNA staining is shown. Scale bar = 10 µm. All presented data are representative of more than three independent experiments.

Immunoblotting of protein extracts of the induced EGFP-RIPK2 cells revealed an upshift of EGFP-RIPK2, starting 2 h post infection (p.i.), and a concomitant decrease of phosphorylation of RIPK2 at S176, whereas NF-κB activation, measured by degradation of IκBα, seemed to start at earlier time points (Fig.1B). Evaluation of the sub-cellular localization of RIPK2 during infection showed that EGFP-RIPK2 localized at F-actin rich *S. flexneri* entry sites at early times of infection (30 min to 1 h p.i.) (Fig. 1C, S1A). However, starting 2 h p.i. formation of EGFP-RIPK2 positive dot-like structures throughout the cell was observed. These structures increased in volume over time, whereas dispersed cytoplasmic localized RIPK2 disappeared (Fig. 1C, Fig. S1C, Movie S1). The appearance of these structures upon infection followed the same kinetic as the change in the electrophoresis migration behavior of RIPK2 (Fig. 1B,C). Staining for nuclear p65 confirmed that NF-κB translocation into the nucleus preceded RIPosome formation (Fig. 1D,E). Staining of *S. flexneri* LPS revealed that RIPosomes did not co-localize with *S. flexneri* (Fig. 1F, S1B). *S. flexneri* infection is known to induce apoptosis in epithelial cells (Carneiro, Travassos et al., 2009, Lembo-Fazio, Nigro et al., 2011). However, the kinetic of activation of caspase-3 upon bacterial infection was not different between EGFP-RIPK2 expressing cells and unmodified HeLa cells (Fig. S2). Furthermore, formation of RIPK2 complexes did not coincide with caspase-3 activation or cell death in single cells (Fig. S2), supporting that RIPK2 complexe formation was not associated with cell death.

Formation of RIPK2 complexes was not limited to *Shigella*, but also seen upon infection with enteropathogenic *Escherichia coli* (EPEC E2348/69). Although EPEC are described as non-invasive pathogens, on epithelial cells, some bacteria can be invasive (Donnenberg, Donohue-Rolfe et al., 1989) (Fig. S3). Infection with EPEC led to RIPosome formation starting from 1 h p.i., whereas EPEC ΔescV, that lack the central component of the type III secretion system (Dupont, Sommer et al., 2016), failed to induce RIPosomes. Visualization of F-actin by Lifeact-Ruby showed pedestal formation at 2 h p.i., which was absent in cells infected with the EPEC ΔescV strain (Fig. S3A and S3D). Similar to *S. flexneri* infection, the amount of RIPosome positive cells after infection with EPEC increased at later time points of infection (Fig. S3C) and was associated with changed migration of the EGFP-RIPK2 protein in SDS-PAGE (Fig. S3E). These data indicate that RIPosome formation is not limited to *S. flexneri* infection, but also occurs during infection with other gram-negative enteropathogenic bacteria that can activate NOD1.

We found EGFP-RIPK2 to be enriched in the triton X-100 insoluble pellet fraction at later times of infection, whereas EGFP-RIPK2 decreased in the triton X-100 soluble fraction upon infection (Fig. 2A). Similar results were obtained for endogenous RIPK2 from HeLa cells infected with *S. flexneri*. Endogenous RIPK2 decreased in the soluble fraction upon infection, whereas it accumulated in the pellet fraction. Though, total RIPK2 levels slightly decreased with infection, an increase of the ratio of the RIPK2 pellet/total fraction at later time points of infection was evident (Fig. 2B), showing that our cell line reflects the properties of the endogenous protein.

**Figure 2:**
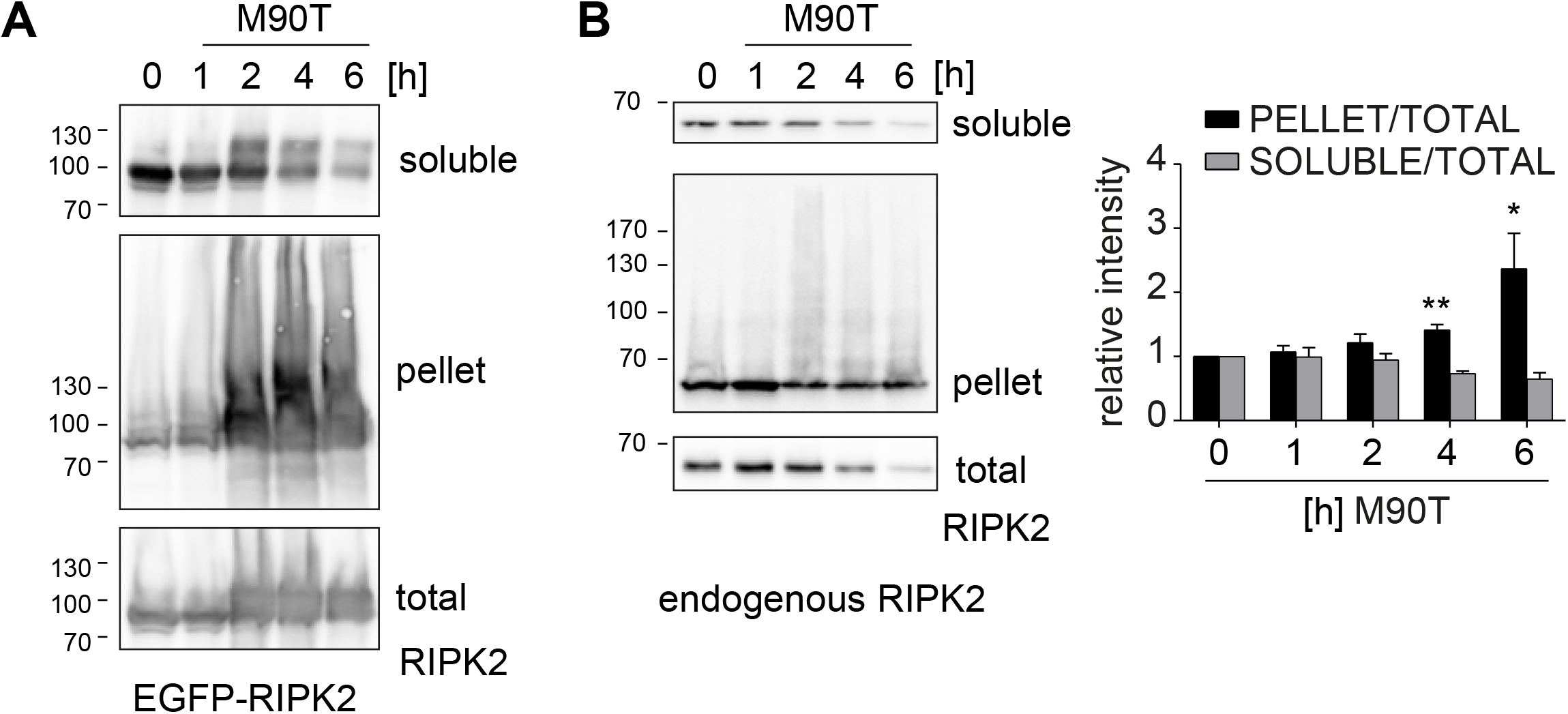
RIPosomes are detergent insoluble. **A:** Analysis of GFP-RIPK2. Whole cell Triton X-100 lysates of HeLa EGFP-RIPK2 cells infected with *S. flexneri* M90T for the indicated time were separated into soluble and insoluble fractions by centrifugation and analyzed by immunoblot. Immunoblot was probed using anti-RIPK2 antibody. **B**: Analysis of endogenous RIPK2. Whole cell Triton X-100 lysates of HeLa cells infected with *S. flexneri* M90T for the indicated time were separated into soluble and insoluble fractions by centrifugation and analyzed by immunoblot. RIPK2 detection was performed using anti-RIPK2 antibody. Right panel: Relative intensity of pellet to total fraction and soluble to total fraction measured by band intensity using ImageJ (n=3). All data are representative of at least two independent experiments. * p<0.05, ** p<0.005 (Students t-test).

To identify if RIPosomes were associated with distinct cellular structures, we next performed co-immunofluorescence, using a set of markers for cellular compartments and innate immune signaling platforms associated with RIPK2. However, RIPosomes did not co-localize with any of the following proteins: TRAF-interacting forkhead-associated protein A (TIFA), which was recently reported to be involved in cytoplasmic innate immune responses towards *Shigella* (Garcia-Weber, Dangeard et al., 2018, Gaudet, Guo et al., 2017), TRAF6 (McCarthy et al., 1998), survival of motor neurons protein protein (SMN), Gemin 3 (Todd, Morse et al., 2010), EEA1 of endosomes (Irving, Mimuro et al., 2014), LC3 from phagosomes (Homer, Kabi et al., 2012), and AIF from mitochondria (Fig. S4).

Taken together, we show that upon activation of epithelial cells by *S. flexneri*, RIPK2 forms distinct cytoplasmic structures. Complex formation followed NF-κB activation and was accompanied by a change in the electrophoretic migration behavior of RIPK2 and formation of triton insoluble aggregates for both endogenous and ectopically expressed RIPK2. RIPK2 complexes did neither co-localize with candidate cellular compartments, signaling platforms nor with bacteria.

### RIPosome formation is dependent on the NOD1/2 CARD

To address whether RIPosome formation upon *Shigella* infection is dependent on NOD1, we performed siRNA-mediated knockdown of NOD1 prior to infection. 2 h p.i. a strongly reduced formation of RIPosomes was observed in NOD1 siRNA treated cells, compared to control siRNA (Fig. 3A). Accordingly, immunoblot analysis showed an upshift of RIPK2 at 6 h p.i. in the control siRNA treated sample, which was virtually absent in the siNOD1 treated sample (Fig. 4B). As expected, NOD1 knockdown reduced IL-8 secretion upon infection, but did not affect IL-8 secretion induced by the overexpression of RIPK2, compared to control siRNA treated cells (Fig. 3C). We further excluded the possibility that RIPosmes were dependent on NF-κB activation. Treatment of the cells with TNF failed to induce RIPosomes (Fig. 3D). Moreover, siRNA targeting of RelA in bacteria infected cells did not change the kinetic of RIPosome formation as indicated by immunoblot and fluorescence microscopy (Fig. 3E). Thus *Shigella*-induced RIPosome formation is an event dependent on NOD1 but independent of NF-κB activation downstream of RIPK2. To analyze the prerequisite for RIPosome formation in greater detail, we used NOD1 and NOD2 with mutations in the NACHT domain, which we previously showed to affect NOD1/2 activation (Zurek, Proell et al., 2012). To this end, NOD1 K208R, D287A, E288A, and H517A and the NOD2 constructs K305R, E383A, and H603A along with NOD1 wt and NOD2 wt as well as the CARD domain of NOD1 alone, were transiently overexpressed in EGFP-RIPK2 cells. All constructs showed membrane localization dependent on the activity of the constructs as reported earlier (Zurek et al., 2012). Surprisingly, all tested NOD1 forms were capable to induce RIPosome formation, except for NOD1 ΔCARD (Fig. 4A). Similarly, all NOD2 mutants were able to induce RIPosomes (Fig. 4B). However, NOD1 and NOD2 were not recruited to RIPosomes in the final stage. Immunoblot analysis confirmed RIPosome formation by upshift of RIPK2 (Fig. 4C), suggesting that the presence of an excess of NOD1/2 CARD is sufficient for RIPosome formation. Taken together, these results showed that RIPosome formation is dependent on the CARD of NOD1/2 but not on NF-κB activation, nor on the possible action of bacterial effectors.

**Figure 3:**
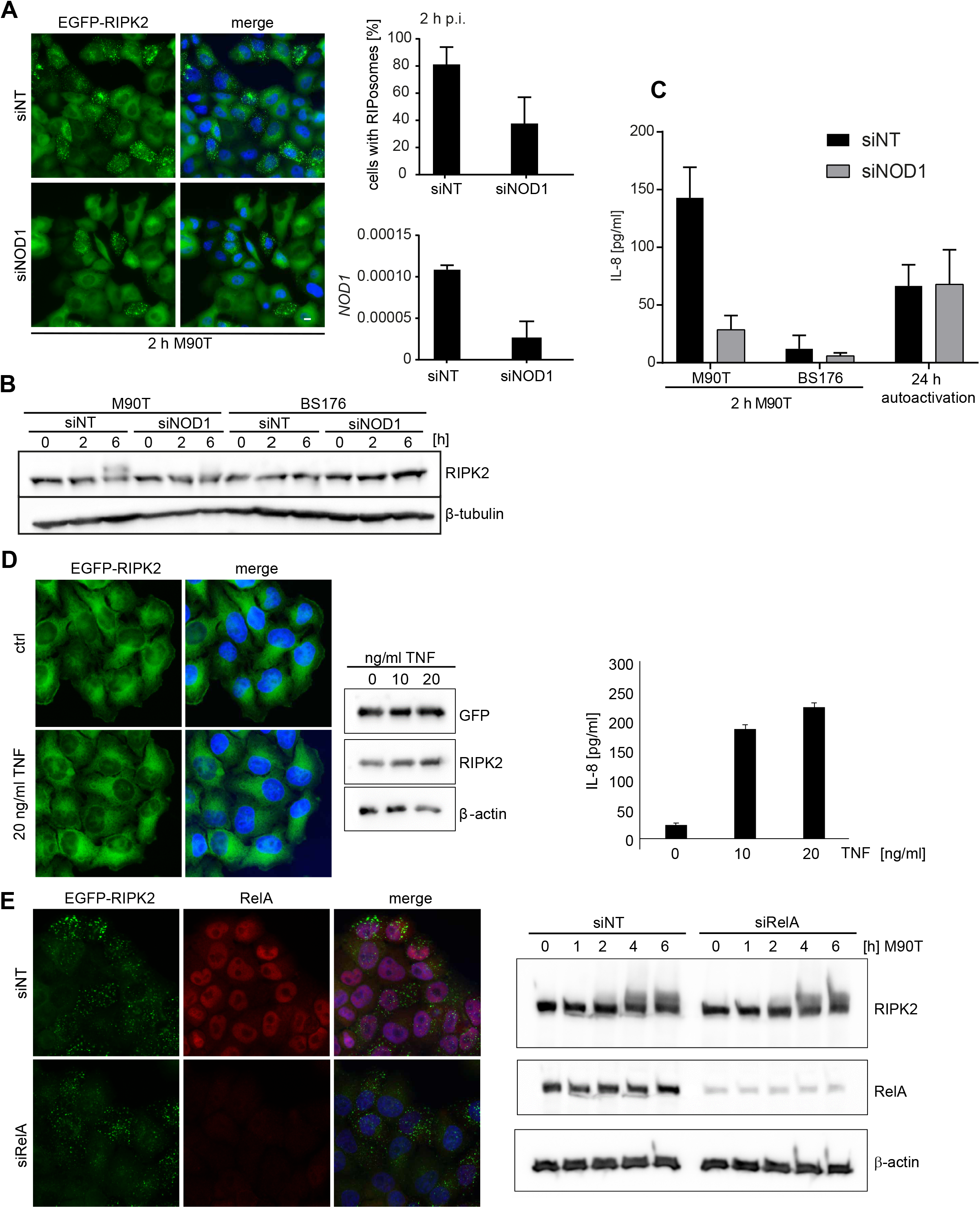
Shigella-induced RIPosome formation depends on NOD1 but not NF-κB activation. **A:** Left panel: Fluorescence micrographs of HeLa EGFP-RIPK2/Lifeact cells, treated for 72 h with *NOD1* siRNA or a non-targeting siRNA, following 2 h *S. flexneri* M90T infection. Doxycycline treatment (1 µg/ml) of the cells was performed 24 h before infection. EGFP-RIPK2 and merge with DNA-staining is shown. Scale bar = 10 µm. Upper right panel: Quantification of RIPosomes in the cells in (A); 500 cells of n=2 experiments were quantified. Lower right panel: RT-qPCR analysis of *NOD1* mRNA expression 72 h after knockdown with a *NOD1* specific or a non-targeting siRNA. n=1, SD of triplicate measurements of one representative experiment is shown. **B:** Immunoblot analysis of HeLa EGFP-RIPK2/Lifeact cells, treated with a *NOD1* specific or a non-targeting siRNA. 24 h before infection, cells were treated with 1 µg/ml doxycycline. Cells were infected 72 h after siRNA treatment with *S. flexneri* M90T or BS176 for the indicated time. Protein levels were detected using anti-RIPK2, and anti-β-tubulin as loading control. **C:** IL-8 ELISA of supernatants from (A) in comparison to supernatants taken 24 h after doxycycline treatment with 1 µg/ml (autoactivation). Mean of n=2 experiments with S.D. is shown. **D:** Left panel: Fluorescence micrographs of HeLa EGFP-RIPK2 cells treated for 18 h with TNF. EGFP-RIPK2 and merge with DNA-staining is shown. Right panel: Immunoblot for GFP-RIPK2 detected by anti-GFP and anti-RIPK2 antibodies in cells treated with different amounts of TNF for 6 h. IL-8 release in the supernatant of these cells is shown in the graph at the right. Mean + S.D. of triplicate measurements of one representative experiment is shown. **E:** HeLa EGFP-RIPK2 cells infected with *S. flexneri* M90T. Left panel: Fluorescence micrographs showing GFP-RIPK2 and RelA staining at 2 h p.i. Right panel: Immunoblot analysis of these cells at different time points post infection.

**Figure 4:**
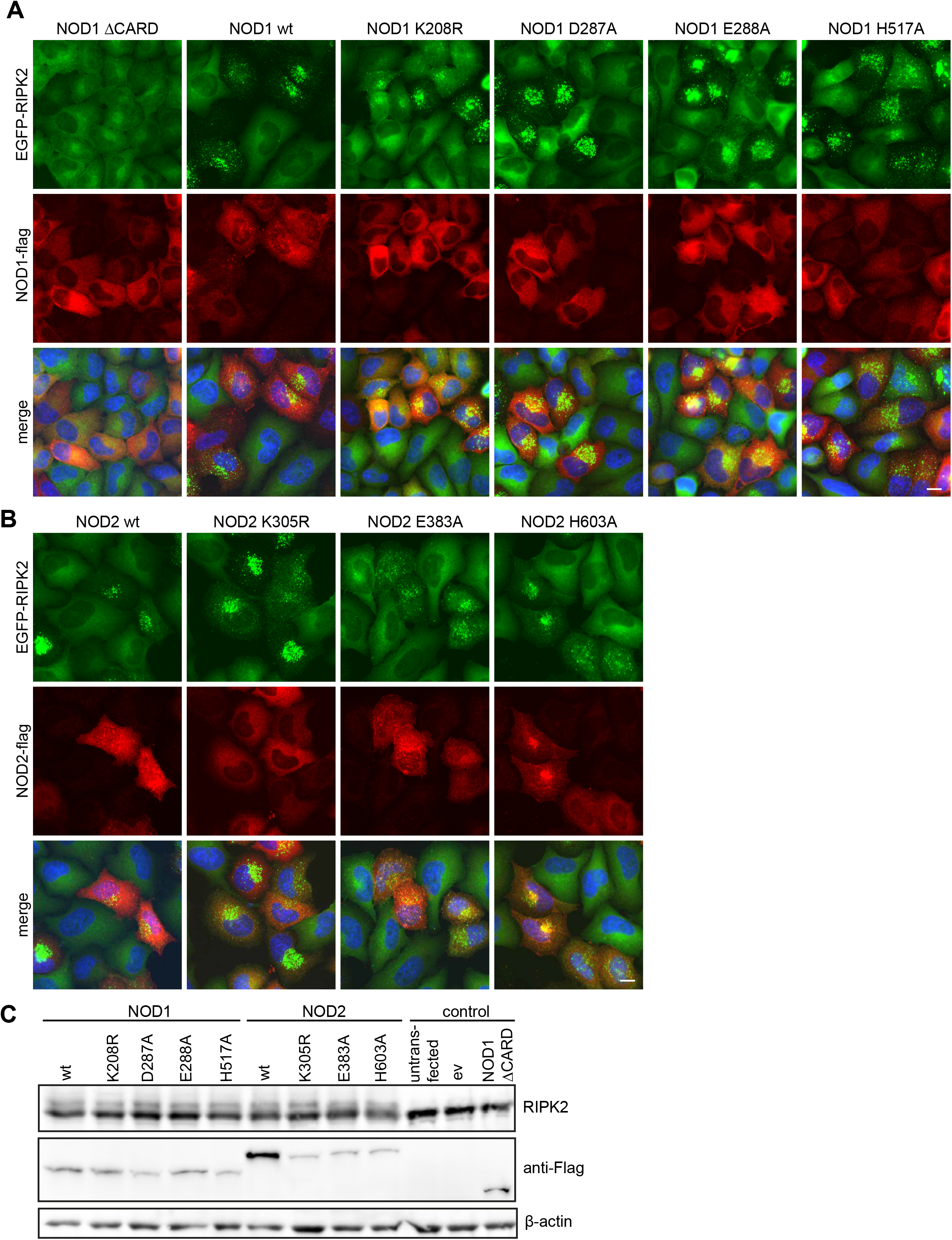
RIPosome formation is dependent on the CARD of NOD1/2. Indirect immunofluorescence micrographs of HeLa EGFP-RIPK2 cells transfected with **(A)** NOD1-Flag or NOD1-Flag mutants or **(B)** NOD2-Flag and or NOD2-Flag mutants. Signals of NOD1/2-Flag, EGFP-RIPK2 and a merge with DNA staining (lower panels) is shown. Scale bar = 10 µm. **C:** Immunoblot analysis of the cells in (A) and (B), probing for RIPK2, Flag and β-actin as loading control.

### XIAP prevents RIPosome formation

Recently, it has been shown that loss of cellular inhibitor of apoptosis proteins (cIAPs) can lead to spontaneous RIPK1/3 Ripoptosome formation, following increased cell death (Feoktistova, Geserick et al., 2011, Tenev, Bianchi et al., 2011). XIAP ubiquitinates RIPK2, a step essential for signaling downstream of NOD1/2 (Krieg et al., 2009). Disruption of the XIAP-RIPK2 interaction thus blocks RIPK2 ubiquitination and NF-κB signaling (Goncharov et al., 2018). Immunostaining of endogenous XIAP in EGFP-RIPK2 cells showed that XIAP was equally distributed throughout the cytoplasm in uninfected cells, but co-localized to RIPosomes at 2 h p.i. (Fig. 5A). siRNA-mediated targeting of XIAP, revealed that knockdown of XIAP was sufficient to induce RIPosome formation, independent of *Shigella* infection (Fig. 5B). In agreement with previous reports, IL-8 secretion, was completely abrogated after XIAP knockdown (Fig. 5C). Immunoblot analysis confirmed upshift of RIPK2 starting 2 h p.i. in cells treated with a control siRNA and infection independent upshift in cells with reduced XIAP levels (Fig. 5D). We noticed that XIAP levels also declined upon bacterial infection and that this coincided with RIPK2 upshift, suggesting that XIAP activity and protein abundance negatively regulate RIPosome formation. Finally, to validate the results obtained by siRNA, we overexpressed second mitochondrial-derived activator of caspases (SMAC), which antagonizes XIAP function (Du, Fang et al., 2000). To this end, ubiquitin fusion constructs, which expose their AVPI motive upon processing in the cytosol were used (Kashkar, Seeger et al., 2006). Both full length and a truncated construct, missing the mitochondrial localization site, induced RIPosome formation, whereas overexpression of the construct missing the AVPI site, which is essential for XIAP antagonization, did not induce RIPosomes nor upshift of RIPK2 in the immunoblot (Fig. 5E). These results demonstrated an essential role of XIAP in preventing RIPosome formation.

**Figure 5:**
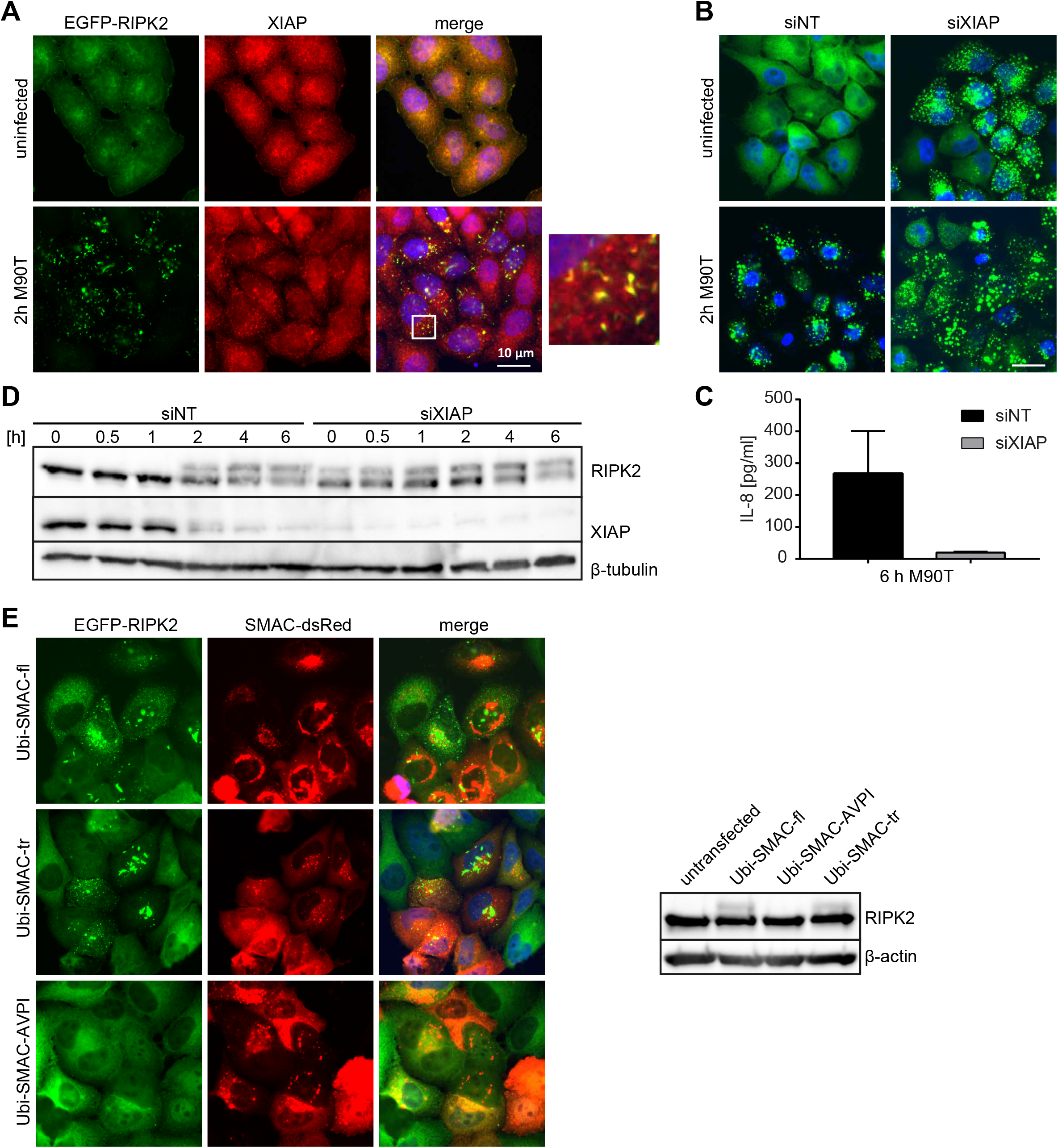
XIAP prevents RIPosome formation. **A:** Indirect immunofluorescence micrographs of HeLa EGFP-RIPK2 cells infected for 2 h with *S. flexneri* M90T. Staining for XIAP, EGFP-RIPK2, and merge with DNA-staining is shown. Co-localization is shown at higher magnification in the inlay. Scale bar = 10 µm. **B:** Fluorescence micrographs of HeLa EGFP-RIPK2 cells, treated for 48 h with a *XIAP* siRNA or a non-targeting siRNA, following infection with *S. flexneri* M90T for 2 h or left uninfected. Cells were treated with 1 µg/ml doxycycline for 24 h prior infection. EGFP-RIPK2 and merge with DNA-staining is shown. Scale bar = 10 µm. **C:** IL-8 release in the supernatants from HeLa EGFP-RIPK2 cells, treated for 48 h with a *XIAP* siRNA or a non-targeting siRNA, following infection with *S. flexneri* M90T for 6 h. Mean of n=2 with S.D. is shown. **D:** Immunoblot analysis of HeLa EGFP-RIPK2 cells infected with *S. flexneri* M90T or BS176 for the indicated time. Immunoblot was probed with anti-RIPK2 antibody, anti-XIAP, and anti-β-tubulin as loading control. **E:** Left panel: Fluorescence micrographs of HeLa EGFP-RIPK2 cells, transfected with 500 ng Ubi-SMAC-fl-pDS-RED, Ubi-SMAC-tr-pDS-RED, or Ubi-SMAC-AVPI-pDS-RED plasmid and treated with 1 µg/ml doxycycline for 24 h. EGFP-RIPK2, SMAC-dsRed, and merge with DNA-staining is shown. Scale bar = 10 µm. Right panel: Immunoblot analysis of cells prepared as in the left panel. Protein levels were detected using an anti-RIPK2 antibody and anti-β-tubulin as loading control.

### Phosphorylation of RIPK2 at Y474 is essential for RIPosome formation

Several autophosphorylation sites of RIPK2 have been reported, S176 and Y474 being the best studied. S176 is described as autophosphorylation site important for RIPK2 catalytic activity (Dorsch et al., 2006), whereas autophosphorylation at Y474 is essential for full NOD signaling (Tigno-Aranjuez et al., 2010). We hypothesized that these phosphorylation sites might have a role in RIPosome formation, as phosphorylation at S176 declined upon infection and subsequent RIPosome formation (Fig. 1B), whereas tyrosine phosphorylation appeared simultaneously (Fig. S5A). Transient overexpression of EGFP-RIPK2 S176A, EGFP-RIPK2 S176E, and EGFP-RIPK2 Y474F in HeLa cells showed that both RIPK2 S176A and RIPK2 S176E induced RIPosomes similar to WT RIPK2, whereas RIPK2 Y474F was not able to form complexes, as shown by immunofluorescence and immunoblot (Fig. S5B,C). In HEK293T cells, expression of increasing amounts of S176A and S176E, but not RIPK2 Y474F, induced NF-κB activation at least as high as RIPK2 wt (Fig. S5D). Stable cell-lines expressing inducible EGFP-RIPK2 S176A, EGFP-RIPK2 S176E, and EGFP-RIPK2 Y474F mutants, confirmed that RIPK2 Y474F was not able to form RIPosomes (Fig. 6A,B).

**Figure 6:**
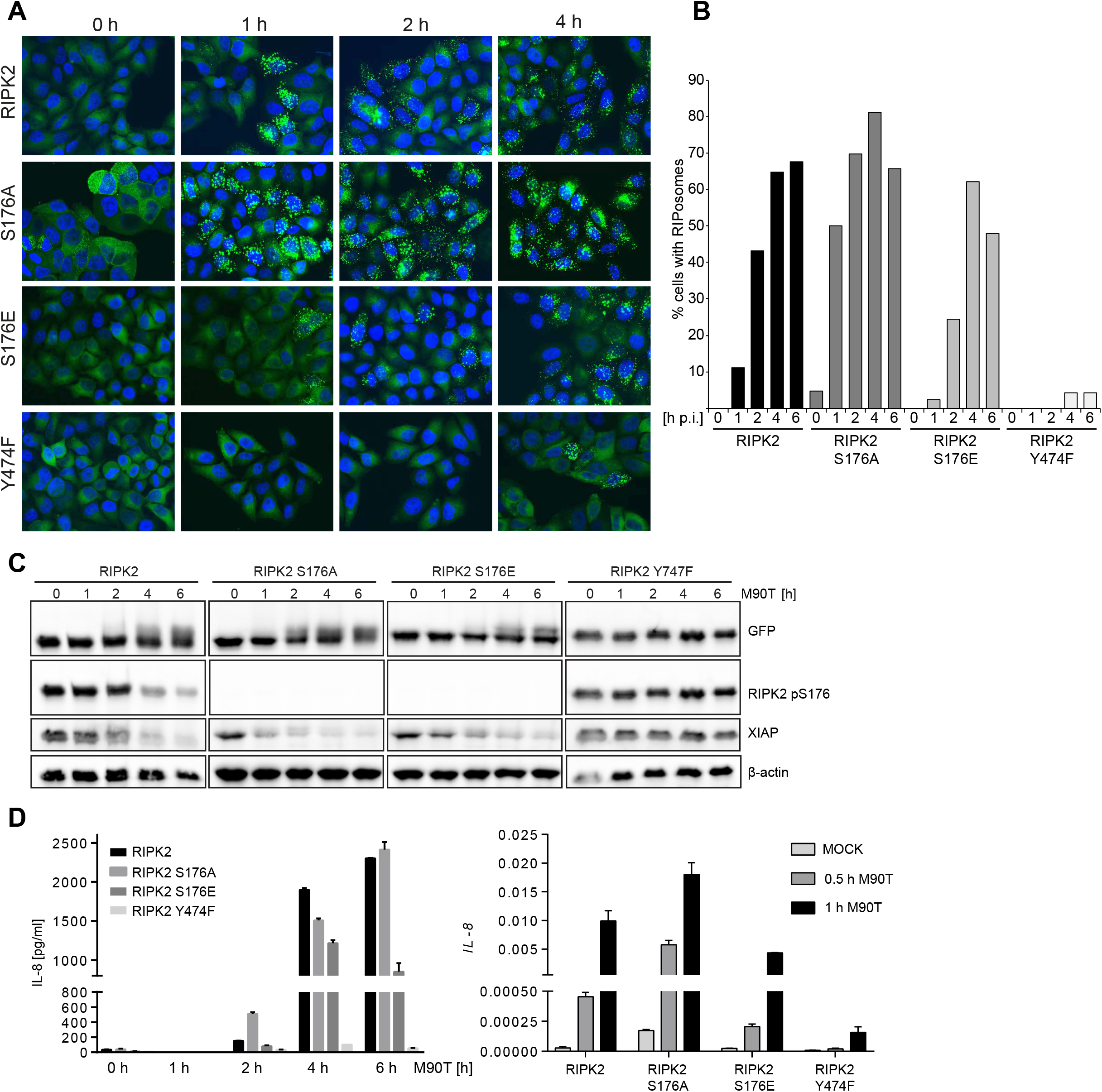
Phosphorylation of RIPK2 at S176 and Y474 contribute to RIPosome formation. **A:** Fluorescence micrographs of stable HeLa lines expressing EGFP-RIPK2, EGFP-RIPK2 S176A, EGFP-RIPK2 S176E and EGFP-RIPK2 Y474F infected for the indicted time with *S. flexneri* M90T. EGFP and merge with DNA-staining is shown. Scale bar = 10 µm. **B:** Quantification of RIPosome containing the cells in (A). At least 180 cell per condition were counted. **C:** Immunoblot analysis of the cells from (A) probing for RIPK2, RIPK2 pS176 and antibody, β-actin as loading control. **D:** Left panel: IL-8 release in the supernatants from the cells in (A,B). Right panel: RT-qPCR of *IL-8* in these cells at early time points of infection.

Using these cell lines with controlled expression of RIPK2, we observed differences in the kinetics of RIPosme formation for the S176 mutants. S176E showed RIPosome formation similar to WT RIPK2 protein, whereas RIPK2 S176A led to RIPosome formation and upshift earlier upon infection, as compared to wt (Fig. 6B,C). Accordingly, RIPK2 S176A induced more IL-8 mRNA expression at early time points of infection, compared to wt RIPK2 (Fig. 6D). By contrast, mutation of Y474 resulted in a RIPK2 protein incapable to induce neither IL-8 nor RIPosomes upon bacterial infection (Fig. 6D). In line with lack of RIPosome formation by RIPK2 Y474F, this protein remained in the soluble fraction upon bacterial infection of the cells (Fig. S5E). Notably, we observed that XIAP levels were reduced upon infection in cells expressing RIPK2, RIPK2 S176A and RIPK2 S176E but not in cells expressing RIPK2 Y747F and correlated with the induction of RIPosome formation (Fig. 6C). As XIAP depletion induced sterile RIPosome formation, this suggest that XIAP might be central in controlling RIPosome formation also upon bacterial infection.

Inhibition of RIPK2 with the tyrosin kinase inhibitor Gefitinib (Tigno-Aranjuez et al., 2010, Tigno-Aranjuez et al., 2014) and the RIPK2 specific compound GSK583 (Haile et al., 2016) (Fig. S6A) also led to spontaneous induction of RIPosomes independent of infection (Fig. S6B). Notably, the appearance of the formed complexes was different for the two compounds: Gefitinib lead to the formation of dot like structures, whereas fiber like structures were obtained with GSK583 (Fig. S6B). RIPosome formation was accompanied by appearance of higher molecular weight signals for RIPK2 and reduced phosphorylation at S176 in infected cells in immunoblot, confirming that this phosphorylation site contributes to RIPosome formation (Fig. S6C). Both substances inhibited *Shigella*-induced IL-8 responses, whereby GSK583 lead to stronger reduction of IL-8 levels compared to Gefitinib (Fig. S6D). However, it was recently shown that these inhibitors interfere with the interaction of RIPK2 with XIAP (Goncharov et al., 2018). This offers a straightforward interpretation of our data as a result of loss of XIAP binding, mimicking depletion or inactivation of XIAP as shown above. In conclusion, these results demonstrate an essential role of phosphorylation of RIPK2 at Y474 for the inflammatory function and RIPosome formation, whereas lack of phosphorylation at S176 can facilitate complex formation.

### Erlin1/2 and 14-3-3 proteins differentially bind to RIPK2

In the next step, we set out to identify cellular proteins, which participate to control RIPK2 functions in the normal and complexed form. To this end, we performed immunoprecipitation of EGFP-RIPK2 from HeLa cells before and at 4 h after infection with *Shigella.* LC-MS/MS analysis of the obtained precipitates led to the identification of several proteins, which were not present in control immunoprecipitates from cells expressing EGFP. Some of these proteins showed differential binding to the soluble and complexed RIPK2. Whereas members of the 14-3-3-family (14-3-3ζ,β,ε,γ) were identified with a high number of peptides in immunoprecipitates prior infection, and the proteins Erlin-1 and Erlin-2 were identified almost exclusively in immunoprecipitates post infection (Fig. 7A). Immunoblot analysis using specific antibodies confirmed that Erlin-1 and Erlin-2 bound to RIPK2 at 4 h p.i. and 6 h p.i., which was concomitant with the upshift of RIPK2, indicative of RIPosome formation, but were not found in the immunoprecipitations from non-infected cells (Fig. 7B). By contrast, binding of 14-3-3ε to RIPK2 was reduced upon infection (Fig. 7B). No change in the stoichiometry of these RIPK2 complexes was observed for RIPK2 Y474F upon infection (Fig. 7B). To validate binding of Erlin-2 to RIPosomes, we performed immunofluorescence analysis. Erlin-2 was distributed throughout the cytosol and localized to nuclear structures in non-infected cells. 4 h p.i. with *Shigella* Erlin-2 localization changed and appeared as dot-like structures in the cytosol, whereas the nuclear staining was lost. These cytosolic Erlin-2 complexes co-localized with RIPosomes (Fig. 7C). Differential binding of Erlin-1 and Erlin-2 to RIPK2 was also observed for the endogenously expressed proteins by immunoprecipitation of RIPK2 from HeLa cells (Fig. 7D), validating the existence of this complex under physiological conditions. As Erlin proteins have been identified as ER lipid raft binding proteins (Browman, Resek et al., 2006, Pearce, Wang et al., 2007, Sprenger, Speijer et al., 2004), we asked if the RIPK2 complexes co-localized to the ER. Immunofluorescence, revealed some, but no strict co-localization of RIPosomes with the ER marker calnexin (Fig. 7E).

**Figure 7:**
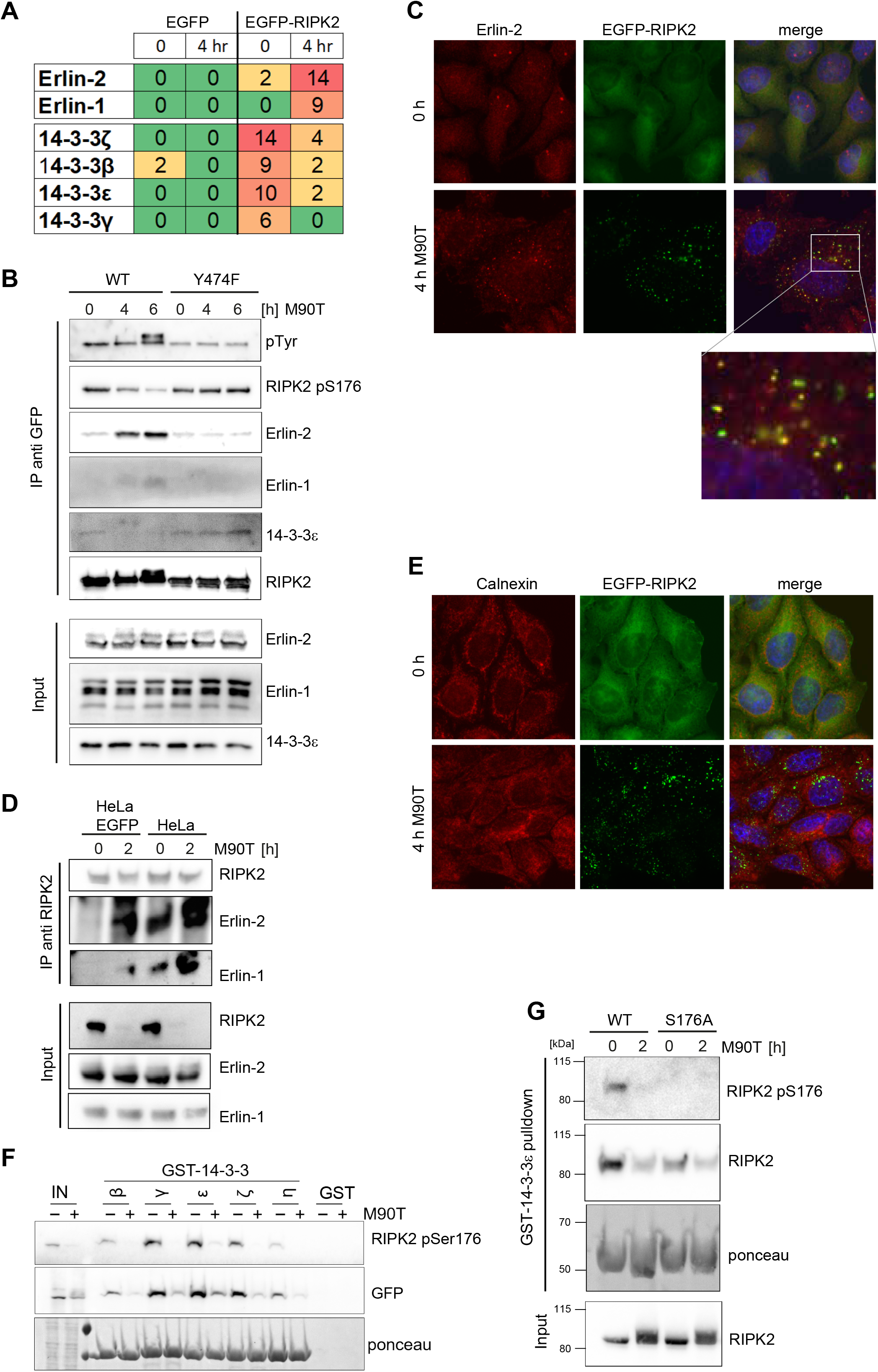
14-3-3 and Erlin proteins differentially bind to RIPosomes. A: Presentation of the proteins with the highest peptide counts identified by MS in the anti-GFP immunoprecipitates from HeLa EGFP-RIPK2 and HeLa EGFP cells used as control. **B:** Immunoblot analysis of anti-EGFP immunoprecipitations from HeLa EGFP-RIPK2 (WT) and HeLa EGFP-RIPK2 Y474F (Y474F) cells infected by *S. flexneri* M90T for 4 h and 6 h. Probing for phosho-thyrosine, RIPK2 pS176 and RIPK2 (RICK) is shown to visualize posttranslational modification of RIPK2 upon infection. Probing for Erlin-1, Erlin-2 and 14-3-3ε confirms differential binding of these proteins to RIPK2 upon infection. **C:** Indirect immunofluorescence micrographs of HeLa EGFP-RIPK2 cells infected for 4 h with *S. flexneri* M90T. Staining for endogenous Erlin-2, EGFP-RIPK2, and merge with DNA-staining is shown. Co-localization is shown at higher magnification in the inlay. **D:** Immunoblot analysis of anti-RIPK2 immunoprecipitations from HeLa cells. Probing for RIPK2 (RICK) and Erlin-1 and Erlin-2 is shown. **E: I**ndirect immunofluorescence micrographs of HeLa EGFP-RIPK2 cells infected for 4 h with *S. flexneri* M90T. Staining for calnexin (ER), EGFP-RIPK2, and merge with DNA-staining is shown. **F:** Immunoblot analysis of pull down experiments using GST-14-3-3 proteins incubated in lysates of HeLa EGFP-RIPK2 cells infected by *S. flexneri* M90T for 2 h. Probing for RIPK2 pS176 and EGFP is shown to visualize co-precipitating RIPK2. Lower panel: Ponceau staining shows equal loads of the GST-14-3-3 proteins. Input fractions are shown in the first two lanes (IN). **G:** Immunoblot analysis of pull down experiments using GST-14-3-3ε protein incubated in lysates of HeLa EGFP-RIPK2 (WT) or HeLa EGFP-RIPK2 S176A (S176A) cells infected by *S. flexneri* M90T for 2 h. Probing for RIPK2 pS176 and EGFP is shown to visualize co-precipitating RIPK2. Lower panel: Ponceau staining to show the 14-3-3 proteins and probing for RIPK2 in the input fractions.

Binding of 14-3-3 proteins to RIPK2 was validated by pull-down assays using recombinant 14-3-3 proteins and lysates from HeLa cells infected with *Shigella*. All tested 14-3-3 isoforms (β,γ,ε,ζ,η) bound RIPK2 in uninfected cells, but only marginally after infection (Fig. 7F). Binding was also reduced, albeit not blocked by mutating S176 to alanine (Fig. 7G), suggesting that other phosphorylation sites might also contribute to 14-3-3 binding.

In summary, we identified novel binding partners of RIPK2, which specifically bind to the different activation forms of RIPK2 and revealed a novel function of Erlin proteins in innate immune processes.

## DISCUSSION

Here, we provide novel insights into the molecular regulation of RIPK2 and show that upon bacterial invasion, RIPK2 forms high molecular weight complexes in the cytosol, which we and others termed RIPosomes (Gong et al., 2018). Formation of these structures was accompanied by reduced detergent solubility of RIPK2 and changes in the electrophoretic mobility. RIPK2 forms hetero-oligomers via CARD:CARD interactions and was found to polymerize to long filamentous structures *in vitro*, which were proposed to act as signaling platforms for the activation of pro-inflammatory signaling downstream of NOD1 and NOD2 (Gong et al., 2018, Pellegrini et al., 2018). Our data support that RIPosome initiation depends on availability of the CARD domain of NOD1/2 that serves as seed for aggregation. Recent *in vitro* structural data that indicate that the CARD of NOD1 forms a transient unstable complex with RIPK2 CARD to induce aggregation is consistent with our *in situ* data (Gong et al., 2018). NF-κB signaling thereby seems to be uncoupled from this process as mutations in NOD1 and NOD2 that interfere with NF-κB activation still triggered RIPosome formation. In addition, mutations in the CARD interaction surface of NOD1 and RIPK2 that interfere with signaling but not with protein interaction are described (Mayle et al., 2014). This suggests that distinct molecular surfaces are involved in signaling versus RIPosome formation. As the NOD1 CARD offers several possible interaction modes with RIPK2, these might define different biological outcomes (Maharana et al., 2017). Formation of higher molecular complexes is also known for other members of the RIPK family. RIPK1 and RIPK3 can form a complex called the Ripoptosome, a cell death platform, discriminating between apoptosis and necroptosis (Feoktistova et al. 2011). However, RIPK2 does not possess the RHIM domain essential for cell death events and oligomerization in RIPK1 and RIPK3. Additionally, our data show that formation of RIPosomes is not directly associated with apoptosis nor cell death in HeLa cells.

Although ectopic expression of RIPK2 was used in some experiemnts, we provide ample evidence to show that this controlled expression of RIPK2 in our stable cell line reflects accurately the endogenous situation. We show that endogenous RIPK2 forms detergent insoluble complexes with similar kinetics and validated interactions of Erlin and 14-3-3 proteins using endogenous proteins. In addition, only the use of this cell lines allowed gaining further insights into RIPK2 activation, as they are so far unique reporters for cytosolic bacterial recognition in real time.

Using this tool, we found that RIPosomes formed after NF-κB activation mediated by NOD1, which is contradictory to a direct role of these structures in NF-κB activation by NOD1/2, as recently proposed by others (Gong et al., 2018, Pellegrini et al., 2018). Activation of NF-κB responses by NOD1 and NOD2 is associated with membrane recruitment of NOD1 and NOD2 (Kufer, Kremmer et al., 2008, Lecine, Esmiol et al., 2007, Nakamura, Lill et al., 2014, Travassos, Carneiro et al., 2010). In line with the formation of RIPosomes after this initial signaling event, we did not observe recruitment of neither NOD1 nor NOD2 to RIPosomes. Moreover, knock-down and inactivation of XIAP, which is essential for NF-κB activation by RIPK2 (Andree, Seeger et al., 2014, Damgaard, Fiil et al., 2013, Goncharov et al., 2018, Krieg et al., 2009), induced RIPosomes in sterile conditions, again supporting that RIPosome formation is independent of inflammatory signaling by RIPK2. *Shigella* induces non-apoptotic SMAC release in epithelial cells to counteract NOD1-mediated sensing by inhibition of XIAP (Andree et al., 2014). This process might be crucial for RIPosome formation upon *Shigella* infection by inhibition of XIAP function. Ubiquitination plays a central role in controlling RIPKs (reviewed in (Witt & Vucic, 2017)). Notably, RIPK1/RIPK3 Ripoptosome formation is also induced by loss of XIAP (Tenev et al., 2011, Yabal, Muller et al., 2014). Mechanistically this was proposed to involve aberrant ubiquitination of RIPK1/3, raising the question which ubiquitination events play a role in RIPK2 complex formation. As RIPK2 is mainly regulated by ubiquitination on K209 via covalent attachment of different linkage-types of ubiquitin (Hasegawa et al., 2008, Panda & Gekara, 2018) this site might play a role in RIPosome formation as well.

The role of XIAP in controlling RIPK2 can also explain why treatment of cells with Gefitinib and GSK583 led to spontaneous formation of RIPosomes as these small compounds interfere with XIAP binding to RIPK2 (Goncharov et al., 2018). XIAP thus might act as a master switch between the NF-κB pathway and RIPosome signaling.

Besides ubiquitination, phosphorylation events play pivotal roles in regulating RIPK2 activity and function. Autophosphorylation of RIPK2 at several sites is associated with activation and dimer formation (Pellegrini et al., 2017). S176 is one of the best-described sites that contributes to RIPK2 activation and functionality (Dorsch et al., 2006, Pellegrini et al., 2017). We observed that RIPosome formation coincided with loss of phosphorylation at this site. Accordingly, mutation of this site to alanine or phosphomimetic glutamic acid did not affect RIPosome formation per se. The S176A mutant showed higher autoactivation and favored complex formation, whereas the phospho-mimetic mutant (S176D) showed decreased activity and delayed complex formation. By contrast, mutation of Y474 to phenylalanine in the CARD completely impaired RIPosome formation and signaling. This is in line with the observation that non-aggregated RIPK2 is present in a highly phosphorylated form (Pellegrini et al., 2018). Phosphorylation at the Y474 corresponding site in ASC was associated with ASC speck formation (Hara, Tsuchiya et al., 2013) and it was proposed that this site in RIPK2 might have similar functions (Boyle, Parkhouse et al., 2014). Our data support a role of this site in aggregate formation for RIPK2. Such phosphorylation patterns might be general regulators of CARD aggregate formation and signaling, as ASC speck formation is dependent on phosphorylation, whereas signaling and interaction of ASC with NLRP3 is not (Hara et al., 2013). In contrast to ASC specks, RIPosomes appeared as multiple high molecular weight complexes per cell and increase over time of infection, suggesting different mechanism of oligomerization of these CARD proteins. The above mentioned phosphorylation events in RIPK2 could also indirectly contribute to RIPosome formation. XIAP protein levels inversely correlated with RIPosome formation for all tested RIPK2 variants and reduction of XIAP levels by siRNA led to spontaneous RIPosome formation. These observations suggest a central role of XIAP in the RIPosome pathway.

Beside XIAP and other E3 ubiquitin ligases, only a few proteins had been identified to bind to RIPK2 and possess regulatory functions. Here, we identified 14-3-3 proteins and Erlin-1 and Erlin-2 (SPFH2) as novel RIPK2 binding proteins. 14-3-3 proteins regulate a plethora of cellular phospho-proteins by interaction with defined phospho-serine/threonine residues. We show that the non-activated soluble form of RIPK2 is a client protein of several 14-3-3 proteins, whereas binding was impaired upon bacterial activation of NOD1-RIPK2. Supporting our results, RIPK2 was identified in immunoprecipitations with 14-3-3 previously (Pozuelo Rubio, Geraghty et al., 2004) and 14-3-3 proteins might regulate RIPK2 signaling as at least 14-3-3-γ appeared as a candidate of a negative regulator of NOD1 and NOD2 signaling in two independent siRNA screens (Bielig, Lautz et al., 2014, Warner, Burberry et al., 2013, Warner, Burberry et al., 2014). Moreover, 14-3-3ζ binds RIPK3 (Zhang, Shao et al., 2009) and regulates its activity by regulating TSC2 binding (Fettweis, Di Valentin et al., 2017). 14-3-3 proteins emerge as important regulators also of immune responses. Innate immune pathways mediated by RIG-I and pyrin, for example, are controlled by 14-3-3 proteins (Gao, Yang et al., 2016, Jeru, Papin et al., 2005, Liu, Loo et al., 2012), and 14-3-3 binding to pyrin has relevant functions in pyrin-associated autoinflammatory disease (Moghaddas, Llamas et al., 2017). Our data add RIPK2 to the list of 14-3-3 client proteins and suggest a role of these proteins in controlling RIPK2-medited innate immune responses. The other protein family identified with high confidence as novel RIPK2 interaction partners were the prohibitin family proteins Erlin-1 and Erlin-2. These localize to ER lipid raft domains (Browman et al., 2006, Pearce et al., 2007, Sprenger, Fontijn et al., 2006) and we observed a partial co-localization of RIPosomes with endoplasmic reticulum (ER), suggesting that Erlin proteins can target RIPK2 complexes to the ER. Erlin-1/2 were also shown to contribute to the ER-associated degradation (ERAD) pathway (Jo, Sguigna et al., 2011, Pearce et al., 2007, Pearce, Wormer et al., 2009, Wang, Pearce et al., 2009). Association of Erlin-2 with detergent resistant membrane domains (DRMDs) is reduced by infection of macrophages with a rough strain of *Brucella melitensis* (Lauer, Iyer et al., 2014). Notably, NOD1/2 and RIPK2 contribute to ER stress induced pro-inflammatory responses upon *Burcella* infection (Keestra-Gounder, Byndloss et al., 2016). This makes it likely that Erlin-2 is involved in regulation of RIPK2-mediated inflammatory signaling and suggests a link of the RIPosome to ER stress response, which is also induced in HeLa cells upon *Shigella* infection via induction of amino acid starvation (Tattoli, Sorbara et al., 2012).

To summarize, our data suggest that RIPosomes are novel signaling platforms, that from in epithelial cells upon bacterial activation of NOD1 and NOD2. These structures might contribute to NF-κB-independent signaling and to limit or terminate RIPK2 activation. The identification of 14-3-3 proteins and Erlin-1 and Erlin-2 as novel binding partners of RIPK2, controlling this cellular process, will help to define novel strategies to target this important pathway in immunity.

## MATERIAL AND METHODS

### Plasmids and reagents

The plasmids pcDNA5/FRT/TO-EGFP and pcDNA5/FRT/TO-EGFP-RIPK2 were generated by molecular cloning. Site directed mutagenesis was used to generate RIPK2-S176A, - S176E and -Y474F. Flag-tagged NOD1, NOD2, and the respective mutants are described in (Kufer, 2008, Kufer, Kremmer et al., 2006). pLifeact-Ruby was kindly provided by Roland Wedlich-Söldner (Riedl, Crevenna et al., 2008). Ubi-SMAC-fl, Ubi-SMAC-tr, and Ubi-SMAC-AVPI were kindly provided by Hamid Kashkar (University of Cologne) (Kashkar et al., 2006). pCR3-Flag-TRAF6 was kindly provided by Greta Guarda (IRB, Bellinzona). pcDNA3_ TIFA-myc was kindly provided by Jessica Yu (Huang, Weng et al., 2012). The pGEX-14-3-3 constructs GST-14-3-3β, GST-14-3-3ϵ, GST-14-3-3γ, GST-14-3-3η and GST-14-3-3σ were obtained from Cheryl Lyn Walker and Judith Bergeron (University of Texas, USA). For transient overexpression, plasmids were transfected, using Lipofectamine2000.

### Cells and cell culture

Stable cell lines were generated by co-transfection of pcDNA5/FRT/TO and pOG44 in a 9:1 ratio into HeLa FlpIN T-REx cells (kindly provided by the Hentze Lab, EMBL, Heidelberg) using Lipofectamine2000 (ThermoFisher Scientific) and selection with 10 µg/ml blasticidin and 500 µg/ml hygromycin B. The HeLa FlpIN T-REx EGFP-RIPK2 Lifeact-Ruby cell line (EGFP-RIPK2/Lifeact) was generated by transfection of pLifeact-Ruby and selection with 250 µg/ml G418. Cell lines were induced with 1 µg/ml doxycycline for 16 h before infection or live cell imaging if not indicated otherwise.

HeLa and HEK293T cells were obtained from ATCC.

Cells were grown in Dulbecco’s modified Eagle’s medium (DMEM) containing 10% heat-inactivated fetal calf serum and 1% penicillin–streptomycin at 37°C with 5% CO_2_. All lines were continuously tested for absence of mycoplasma by PCR.

### Bacteria and bacterial infection

*Shigella flexneri* M90T *afaE* and BS176 *afaE* (Clerc & Sansonetti, 1987) were kindly provided by Philippe Sansonetti (Institute Pasteur, Paris) and were grown in Caso Broth containing 200 µg/ml Spectinomycin. EPEC E2348/69 wt (Streptomycin^R^) and EPEC E2348/69 δescV (escV::miniTn10*kan*, Streptomycin^R^, Kanamycin^R^) were kindly provided by Mathias Hornef (RWTH Aachen) were and grown in Caso Broth or LB supplemented with appropriate antibiotics (Dupont et al., 2016).

Infection with *Shigella* was performed at a MOI of 10 in DMEM without supplements. After 15 min of bacterial sedimentation at room temperature, infection was started at 37°C and 5% CO_2_ After 30 min the medium was changed to DMEM containing 100 µg/ml gentamycin. For ELISA, supernatants were collected 2 h or 6 h post infection.

EPEC infection was performed in 24-well plates at a MOI of 25 in a volume of 250 µl DMEM without supplements. EPEC were centrifuged onto the cells for 5 min at 560 x g, followed by infection at 37°C and 5 % CO_2_ For gentamycin protection assay, cells were lysed with 1 ml 0.1% Triton-X-100 in PBS, serial dilutions were prepared in PBS, streaked onto agar plates containing 200 µg/ml spectinomycin, and incubated overnight at 37°C. The next day, colony formation was analyzed.

### siRNA knockdown and gene expression analysis

Knockdown of XIAP and NOD1 was performed using Hiperfect transfection (QIAGEN) of the following siRNAs: siXIAP (AAGGAATAAATTGTTCCATGC, QIAGEN), siNOD1 (Hs_ CARD4_ 4, SI00084483, QIAGEN), siRelA (AAGATCAATGGCTACACAGGA, QIAGEN) and a non-targeting siRNA (All-Star negative control, QIAGEN).

For gene expression analysis RT-qPCR analysis was performed. 2 µg of total RNA was transcribed into cDNA, using the RevertAid First Strand cDNA Synthesis Kit (ThermoFisher Scientific). qPCR was performed using iQ SYBR Green Supermix (BioRad). The following primers were used: NOD1_fwd: TCCAAAGCCAAACAGAAACTC, NOD1_ rev: CAGCATCCAGATGAACGTG; GAPDH_ fwd: GGTATCGTGGAAGGACTCATGAC, GAPDH_ rev: ATGCCAGTGAGCTTCCCGTTCAG; IL-8_ fwd: ATGACTTCCAAGCTGGCCGTGGCT, IL-8_ rev: TCTCAGCCCTCTTCAAAAACTTCTC.

### Fractionation, GST-14-3-3 pulldown assays, immunoprecipitation and MS analysis

Native cell lysates were prepared by solubilizing cells in Triton-X-100 lysis buffer (150 mM NaCl, 50 mM Tris pH 7.5, 1% Triton-X-100) including proteinase and phosphatase inhibitors (1x Roche complete mini pill, 20 µM β-Glycerophosphate, 100 µM Sodium-Orthovanadate, 5 mM Sodium fluoride) followed by centrifugation for 10 min at 4°C with 21.130 x g to separate soluble lysate and insoluble fractions.

Recombinant GST-14-3-3 proteins were produced in *E.coli* and purified as previously described (Hausser, Link et al., 2006). Pulldowns were performed by incubating equal amounts of protein lysates with GST-14-3-3 coupled to glutathione sepharose beads for 3 hours at 4°C. Beads were washed three times with lysis buffer. For immunoprecipitation of endogenous RIPK2 equal amounts of cell lysates were incubated with 2 µg specific antibodies at 4°C over night. Immune complexes were collected with Affi-prep Protein A resin (Bio-Rad) and washed three times with lysis buffer. For immunoprecipitation of EGFP-RIPK2 equal amounts of cell lysates were incubated with GFP-Trap_A beads (Chromotek) and washed three times with lysis buffer. Samples for SDS-PAGE were denatured by boiling in SDS buffer (Laemmli).

### NanoLC-MS/MS analysis

Proteins were digested on beads using trypsin (Roche, Germany) in 6 M Urea, 50 mM Tris-HCl pH 8.5. Cysteines were reduced using DTT and then alkylated by Chloroacetamide. Samples were then diluted to a final concentration of 2 M Urea. 750 ng trypsin were added and samples were digested overnight at 25 °C. The digests were stopped by adding trifluoroacetic acid (TFA). Next, peptide mixtures were concentrated and desalted on C18 stage tips and dried under vacuum. Samples were dissolved in 0.1 % TFA and were subjected to nanoLC-MS/MS analysis on an EASY-nLC 1000 system (Thermo Fisher Scientific, Germany) coupled to a Q-Exactive Plus mass spectrometer (Thermo Fisher Scientific, Germany) using an EASY-Spray nanoelectrospray ion source (Thermo Fisher Scientific, Germany). The system was controlled by Xcalibur 3.0.63 software. Tryptic peptides were directly injected to an EASY-Spray analytical column (2 μm, 100 Å PepMapRSLC C18, 25 cm (UniProt Consortium, 2018)10 min, 15 min isocratic at 90% solvent B (solvent A: 0.5% acetic acid;, solvent B: in ACN/H_2_O (80/20, v/v), 0.5% acetic acid).

MS spectra (m/z = 300-1600) were detected at a resolution of 70000 (m/z = 200) using a maximum injection time (MIT) of 100 ms and an automatic gain control (AGC) value of 1 × 10^6^ MS/MS spectra were generated for the 10 most abundant peptide precursors using high energy collision dissociation (HCD) fragmentation at a resolution of 17500, normalized collision energy of 27 and an intensity threshold of 1.3 × 10^5^. Only ions with charge states from +2 to +5 were selected for fragmentation using an isolation width of 1.6 Da. For each MS/MS scan, the AGC was set at 5 × 10^5^ and the MIT was 100 ms.

Mascot 2.6 (Matrix Science, UK) was used as search engine for protein identification. Spectra were searched against a human UniProt database (UniProt Consortium, 2018). Scaffold 4.8.6. (Proteome Software, USA) was used to evaluate peptide identifications. These were accepted with a peptide probability greater than 80.0% as specified by the Peptide Prophet algorithm (Keller, Nesvizhskii et al., 2002). Proteins had to be identified by at least two peptides and a protein probability of at least 99% to be accepted.

### Immunoblotting

Cells were lysed in SDS buffer (Laemmli), followed by protein denaturation for 5 min at 95°C. Proteins were separated by SDS-PAGE and blotted to 0.2 µm nitrocellulose membrane, followed by blocking with 0.5 % Roche Blocking in PBS and incubation with the appropriate antibodies. As primary antibodies anti-RIPK2 (A-10, sc-166765, Santa Cruz), anti-RIPK2 (H-300, sc-22763, Santa Cruz), anti-IκBα (#4814, Cell Signaling), anti-pEIF2α (#3398, Cell Signaling), anti-EIF2α (#5324, Cell Signaling), anti-Flag (Sigma, F1804), anti-β-tubulin (T7816, Sigma), anti-β-actin (sc-47778, Santa Cruz), anti-XIAP (E-2, sc-55551, Santa Cruz), anti-Erlin-1 (sc-514820, Santa Cruz), anti Erlin-2 (ABS1610, Sigma), anti-14-3-3ε (F-3, sc-393177, Santa Cruz) were used. For detection of primary antibodies anti-rabbit-HRP (170-6515, BioRad), and anti-mouse-HRP (170-6516, BioRad) were used. Detection of the signals were performed using Clarity Western ECL Substrate (BioRad), or SuperSignal West Femto maximum sensitivity substrate (ThermoFisher Scientific) on an automated high sensitivity camera system (Fusion FX, VilbertLourmat).

### Immunofluorescence microscopy

For immunofluorescence, cells were fixed using 4% paraformaldehyde in phosphate-buffered saline (PBS) for 15 min at room temperature, following permeabilization with 0.1% Triton X-100 for 5 min. Blocking was performed using 5% fetal calf serum in PBS, following incubation with the primary antibody over night at 4°C. Primary antibodies used were anti-*Shigella*-LPS (rabbit anti-*Shigella flexneri* LPS 5a, kindly provided by the lab of Philippe Sansonetti, Institute Pasteur Paris), anti-p65 (sc-8008, Santa Cruz), anti-cleaved-caspase-3 (Asp175, #9661S, Cell Signaling), anti-XIAP (M044-3, MBL), anti-SMN (sc-32313, Santa Cruz), anti-Gemin3 (sc-57007, Santa Cruz), anti-EEA1 (sc-53939, Santa Cruz), anti-LC3β (NEB, 3868), anti-calnexin (#2433, Cell Signaling), Anti-Erlin-2 (ABS1610, Sigma), anti-Flag M2 (F1804, Sigma). After washing in PBS, incubation with the secondary antibody was performed for 1 h at room temperature. The secondary antibodies used were Alexa 546-conjugated goat anti-rabbit IgG, and Alexa 546-conjugated goat anti-mouse IgG (Molecular Probes). Coverslips were mounted in Mowiol-Hoechst (Hoechst 33258, Sigma).

For Live Cell Imaging, cells were cultured in glass bottom petri dishes in FluoroBrite DMEM (ThermoFisher Scientific), induced with doxycycline (1 µg/ml) for 16 h and imaged at 37°C in a 5 % CO_2_ atmosphere. Imaging was performed using an automated Leica DMi8 microscope with incubation chamber and the HC PL APO x63/1.40 oil objective. Images were processed using the Leica LasX software.

### Quantification of RIPosomes

RIPosomes were quantified by eye using blinded processed images. Cells with three or more small bright dots were counted as RIPosome positive cells. At least 200 cells from one experiment were counted for p65 nuclear translocation, 500 cells for EPEC infection kinetics, and 500 cells for *NOD1* knockdown from two independent experiments.

### NF-κB luciferase reporter assays

NF-κB activation was measured by a luciferase-reporter gene assay. 30 000 cellsl were plated per well of a 96-well plate and transiently transfected with X-treme GENE9. 8.6 ng of β-galactosidase plasmid, 13 ng of the luciferase-reporter-plasmid and 10 ng, 5 ng or 1 ng of EGFP-RIP2-expression plasmids were transfected into the cells, using a total DNA amount of 51 ng plasmid per well. After incubation, cells were lysed and luciferase activity quantified on a luminometer. β-galactosidase activity as measured by ONPG assay. Luciferase activity was normalised to the β-galactosidase activity.

### ELISA

IL-8 (CXCL8) was measured in cell culture supernatants using a Duoset (DY208, Bio-Techne) according to the manufacturer’s instructions.

### Statistical analysis

Data was analyzed and plotted using Microsoft Excel and GraphPad Prism 7.0.

## ACKNOWLEDGEMENTS

We thank Berit Würtz (MS Core facility of the University of Hohenheim) for help with the LC/MS-MS analysis and Mariana Mohr for help in generating the EGFP-RIPK2 line. This work was supported by the German research foundation (DFG) grant KU 1945/4-1 to TAK.

## AUTHOR CONTRIBUTIONS

CA, KE, SB and IK performed the experiments.

JP performed MS analysis and data processing.

TAK conceived and supervised the study.

CA, KE and TAK wrote and edited the manuscript.

## Supplemental Information

**Movie S1: Formation of RIPosomes in HeLa EGFP-RIPk2 cells upon Shigella infection.**

**Fig. S1.**
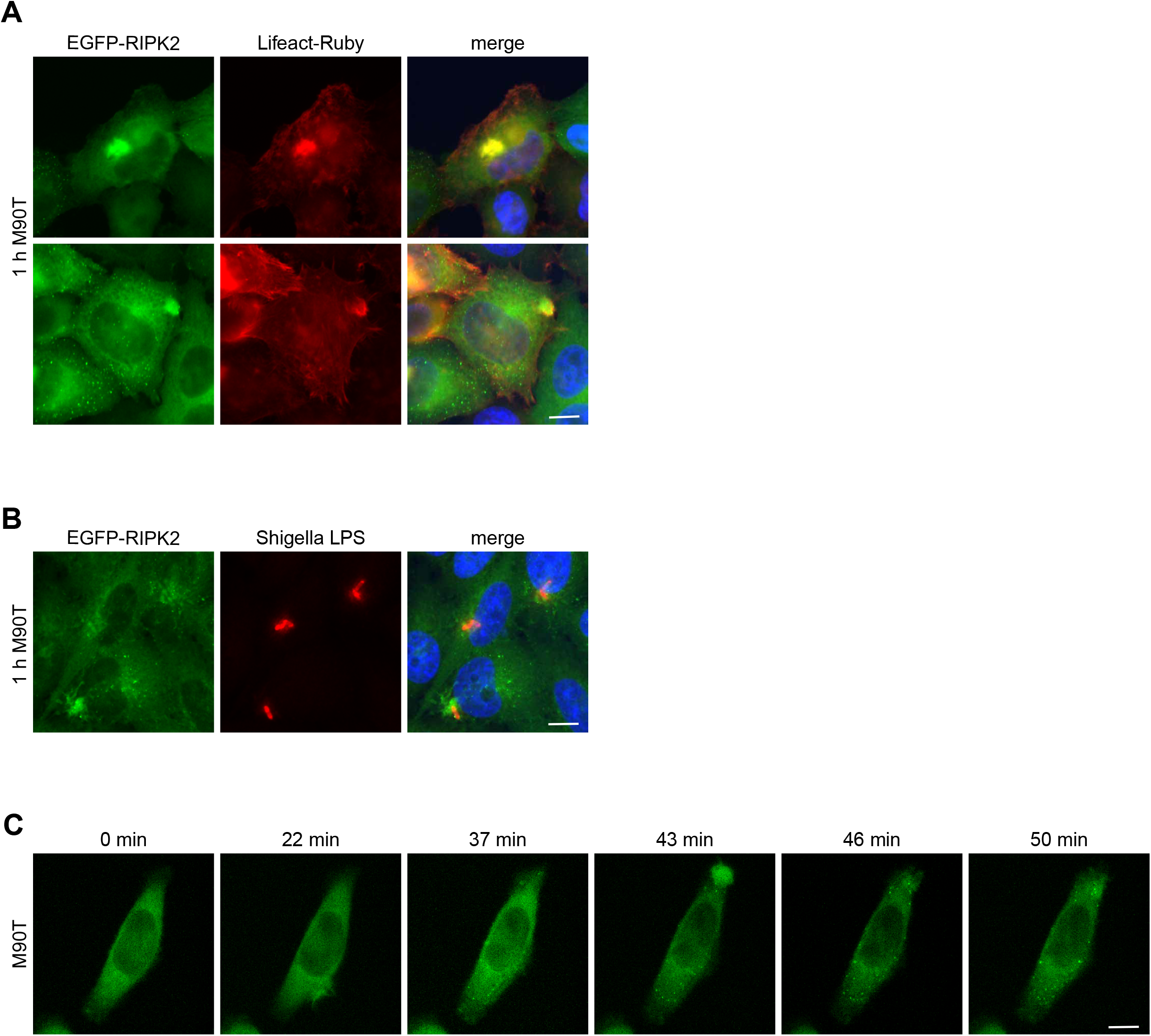
Cellular localization of RIPK2 at early times of infection with *Shigella*. **A:** Fluorescence micrographs of HeLa EGFP-RIPK2/Lifeact cells infected with *S. flexneri* M90T for 1 h. EGFP-RIPK2, Lifeact-Ruby, and merge with DNA-staining is shown. Scale bar=10 µm. **B:** Indirect immunofluorescence micrographs of HeLa EGFP-RIPK2 cells infected with *S. flexneri* M90T infection for 1 h. Immunostaining was performed using an anti-*Shigella*-LPS antibody together with DNA-staining. Scale bar=10 µm. **C:** Single images from movie S1 of HeLa EGFP-RIPK2/Lifeact cells, treated with 1 µg/ml doxycycline for 16 h, following *S. flexneri* M90T infection. EGFP-RIPK2 is shown. Scale bar=10 µm.

**Fig. S2.**
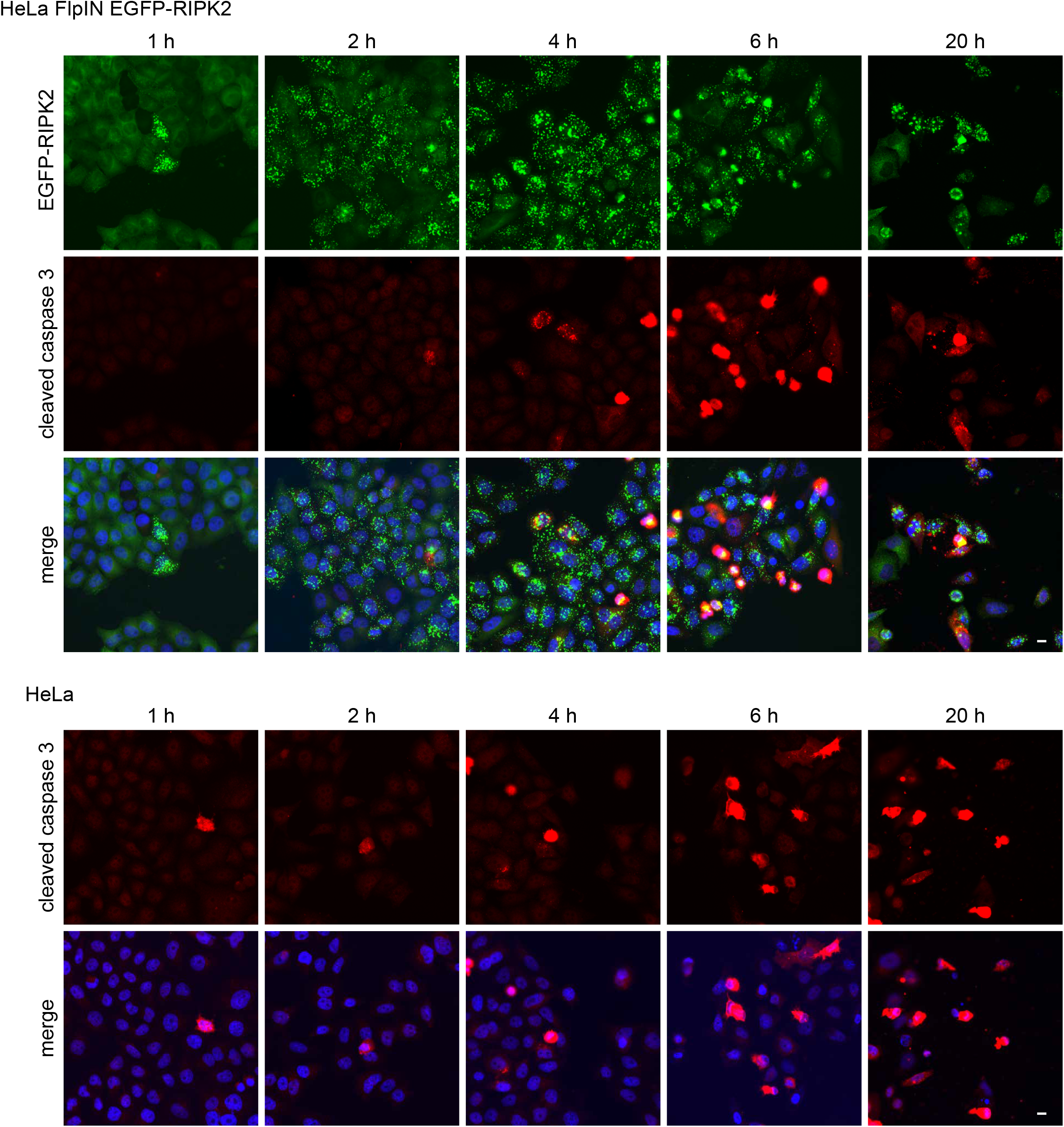
RIPosome formation is not linked to apoptosis. Indirect immunofluorescence micrographs of *S. flexneri* M90T infected HeLa EGFP-RIPK2 cells (upper panel) and HeLa cells (lower panel) immunostained for cleaved caspase 3. HeLa EGFP-RIPK2 cells infected with *S. flexneri* M90T for the indicated time. Signal of EGFP and anti-cleaved caspase 3 antibody and a merge with DNA-staining is shown. Scale bar =10 µm.

**Fig. S3.**
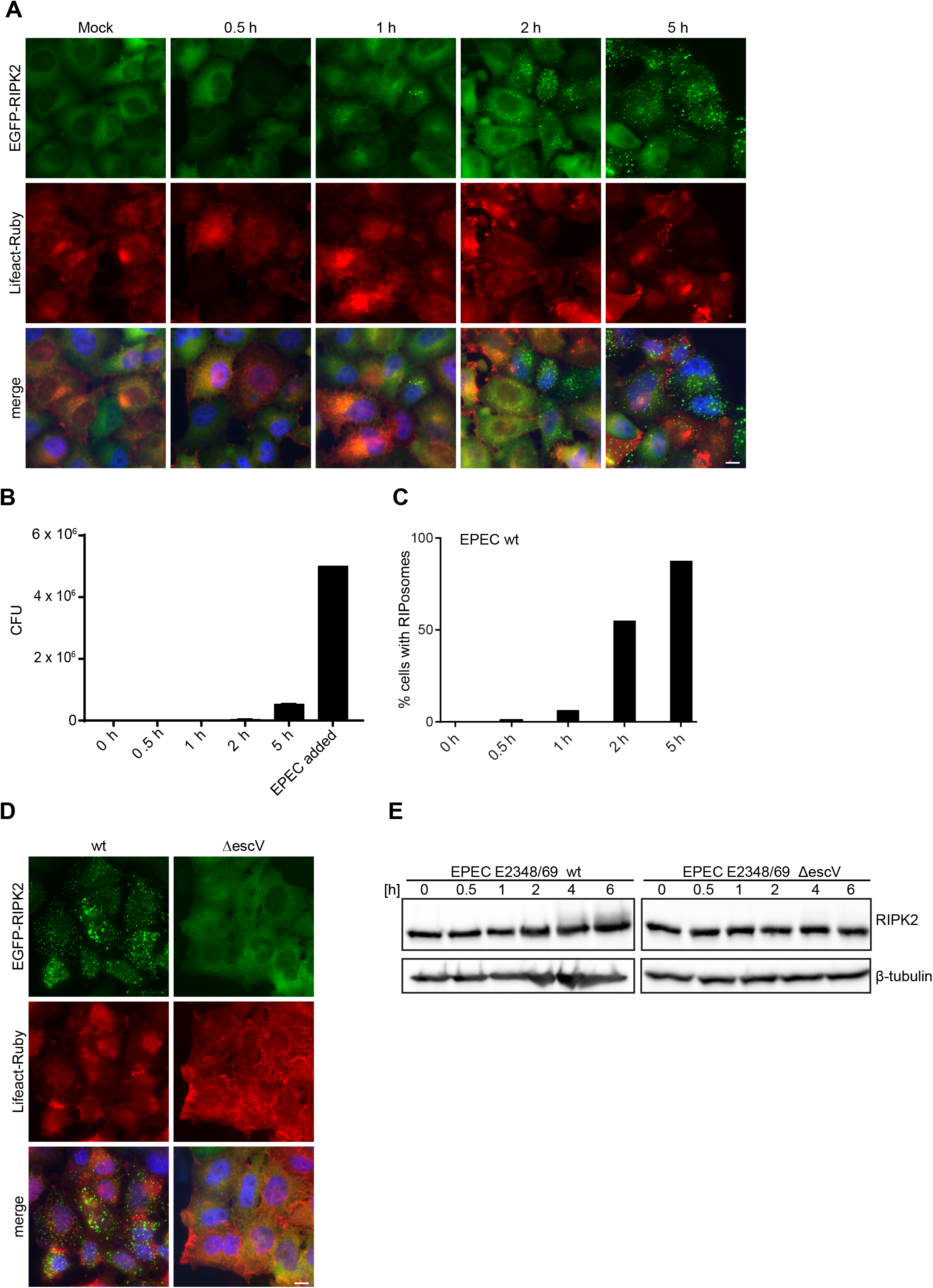
EPEC infection leads to RIPosome formation. **A:** Fluorescence micrographs of HeLa EGFP-RIPK2/Lifeact cells infected with EPEC E2348/69 wt. Gentamycin (final concentration: 100 µg/ml) was added 0 h (directly after centrifugation), 0.5 h, 1 h, or 2 h after infection and incubated for a total infection time of 5 h. Infection for 5 h was performed without gentamycin. EGFP-RIPK2, Lifeact-Ruby, and merge with DNA-staining is shown. Pedestal formation is visible for cells with attached bacteria. Scale bar = 10 µm. **B:** Gentamycin protection assay of HeLa EGFP-RIPK2 Lifeact-Ruby cells, infected with EPEC E2348/69, following medium change to gentamycin containing medium (100 µg/ml) at indicated time points. Cells were incubated for a total time of 5 h. **C:** Quantification of RIPosomes. In total 500 cells of **(A)** were quantified. **D:** Fluorescence micrographs of HeLa EGFP-RIPK2/Lifeact cells infected with EPEC E2348/69 or EPEC E2348/69 ΔescV for 6 h. EGFP-RIPK2, Lifeact-Ruby, and merge with DNA-staining is shown. Scale bar = 10 µm. **E:** Immunoblot analysis of HeLa EGFP-RIPK2 cells infected with EPEC E2348/69 or EPEC E2348/69 ΔescV for the indicated time. 2 h post infection the medium was changed to medium containing 100 µg/ml gentamycin. Immunoblot was probed with anti-RIPK2, and anti-β-tubulin as loading control.

**Fig. S4.**
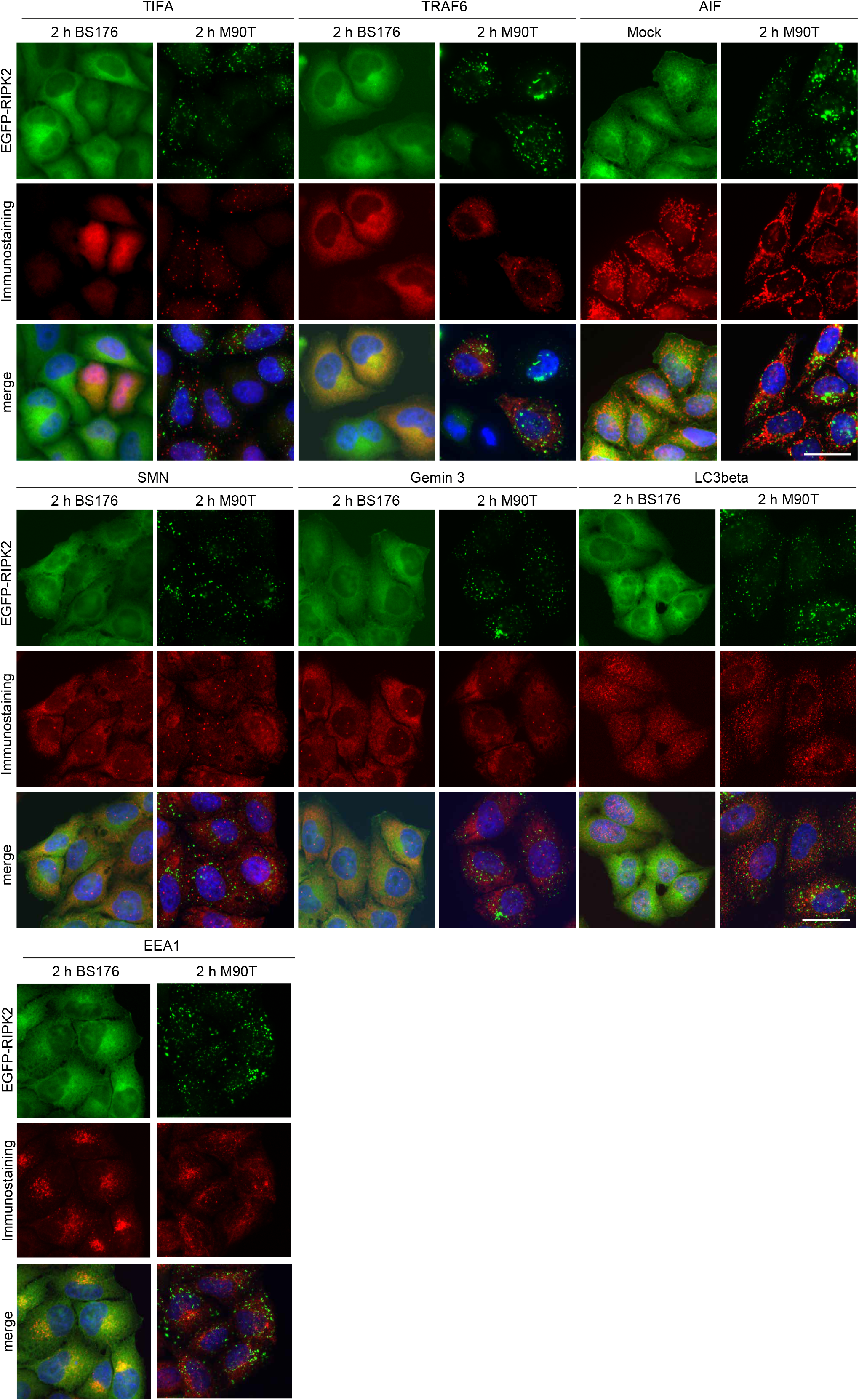
RIPosomes co-localization with cellular compartments and signaling platforms. Indirect immunofluorescence micrographs of HeLa EGFP-RIPK2 cells infected with *S. flexneri* M90T or BS176 for 2 h. TIFA-Flag and Flag-TRAF6 were transfected 16 h before doxycycline induction. Immunostaining of cellular markers was performed using: anti-Flag, anti-AIF, anti-SMN, anti-Gemin3, anti-EEA1, anti-LC3β antibodies together with DNA-staining. Scale bar = 10 µm.

**Fig. S5.**
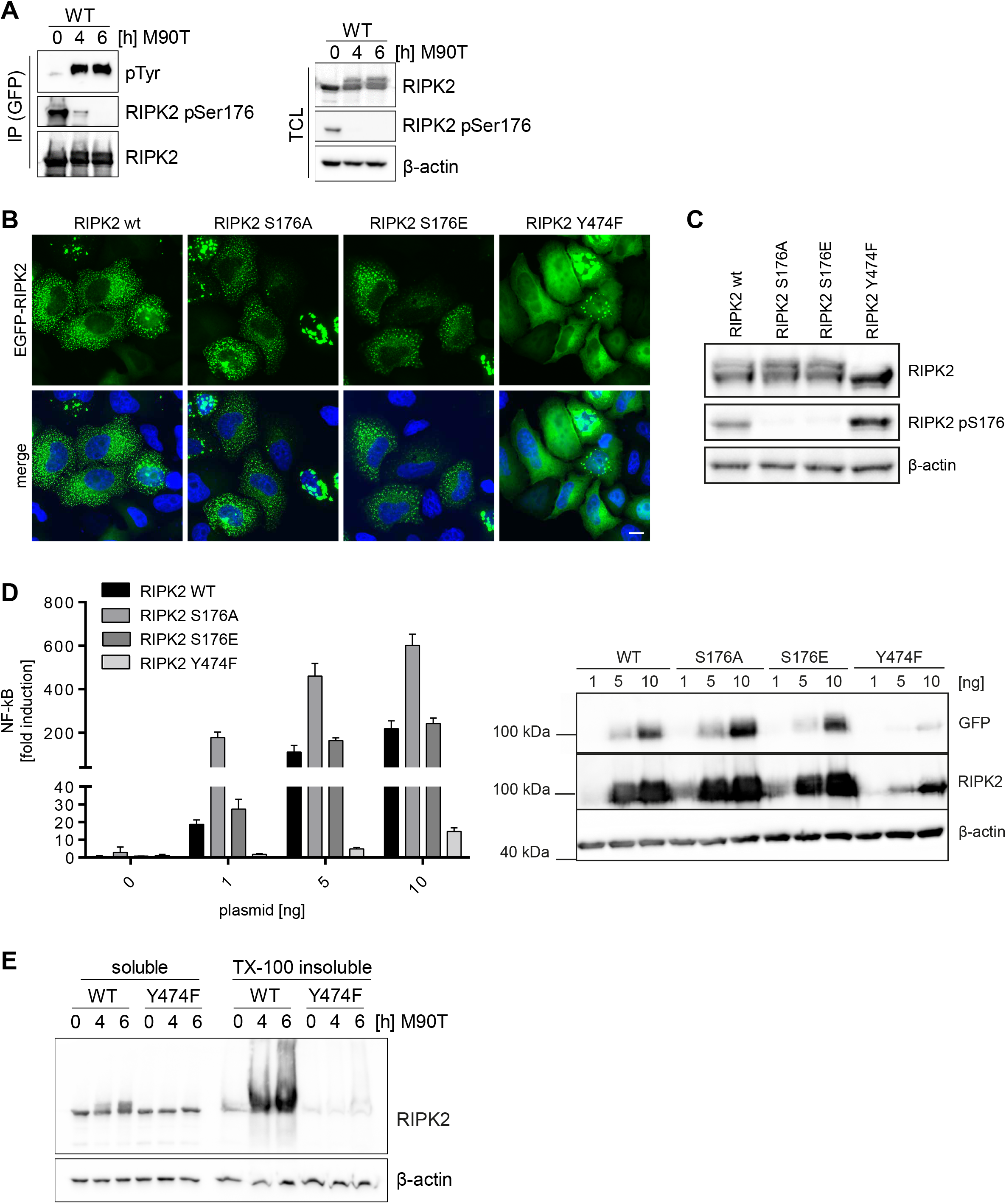
Effect of the mutation of RIPK2 phosphorylation sites. **A:** Immunoblot analysis of anti-GFP immunoprecipitates of EGFP-RIPK2 cells following infection with *S. flexneri* M90T. Probing for RIPK2, RIPK2 pS176, pTyr and β-actin as loading control. Immunoprecipiated proteins are shown in the left panel (IP) and input fraction (total cell lysate, TCL) in the right panel. **B:** Fluorescence micrographs of HeLa EGFP-RIPK2 cells transiently transfected with EGFP-RIPK2, EGFP-RIPK2 S176A, EGFP-RIPK2 S176E or EGFP-RIPK2 Y747F. EGFP and merge with DNA-staining is shown. Scale bar = 10 µm. **C:** Immunoblot analysis of TCL from the cells from (B). Probing for RIPK2, RIPK2 pS176 and β-actin as loading control is shown. **D:** NF-κB luciferase reporter assay in HEK293T cells transfected with the indicted amounts of EGFP-RIPK2, EGFP-RIPK2 S176A, EGFP-RIPK2 S176E or EGFP-RIPK2 Y747F. Right panel: Expression of the respective proteins was controlled by immunoblot probing for GFP, RIPK2 and β-actin as loading control. **E:** Whole cell Triton X-100 lysates of HeLa EGFP-RIPK2 and EGFP-RIPK2 Y747F cells infected with *S. flexneri* M90T for the indicated time were separated into soluble and insoluble fractions by centrifugation and analyzed by immunoblot. Immunoblot was probed for RIPK2 and β-actin as loading control.

**Fig. S6.**
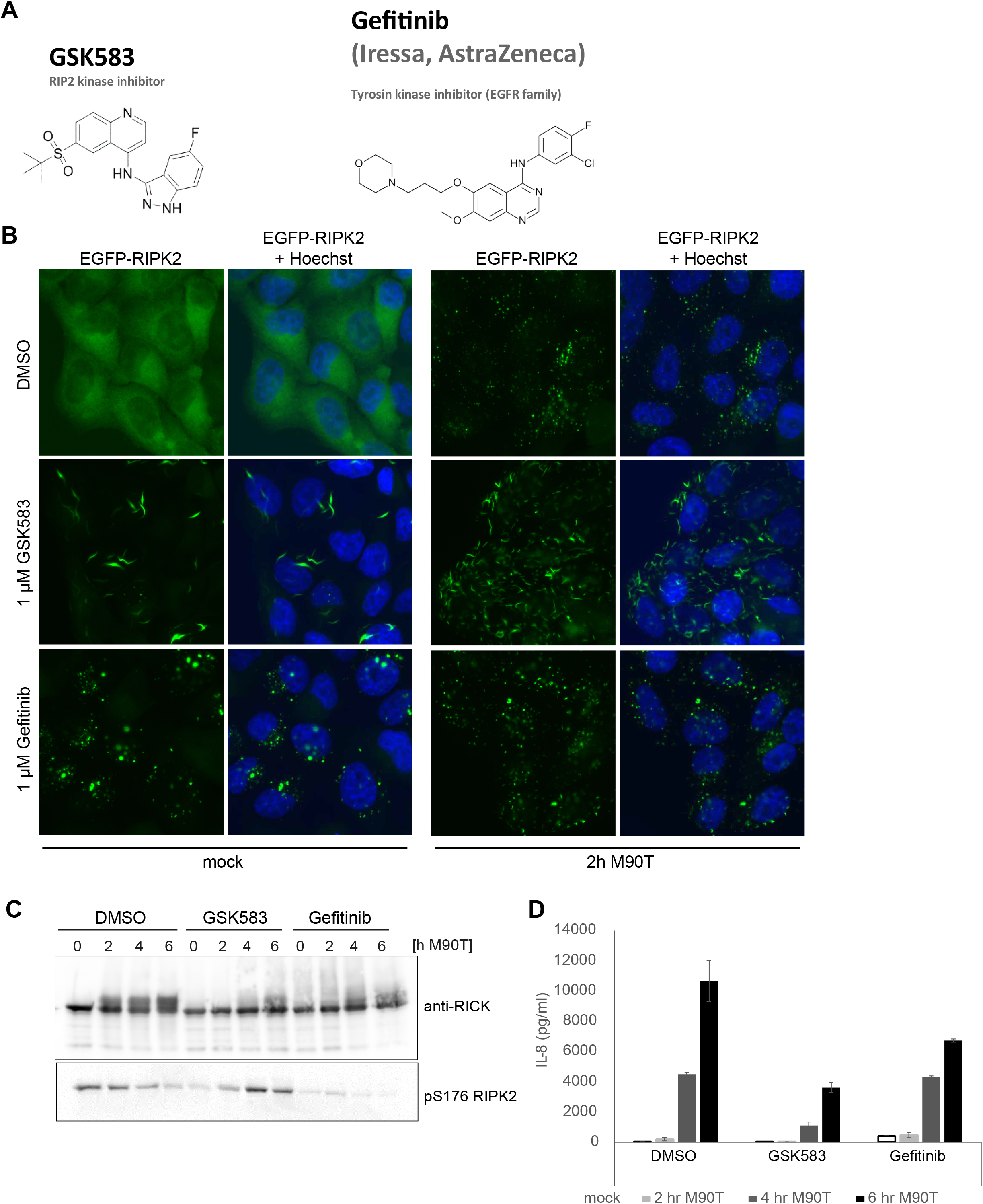
Effect of RIPK2 inhibitors on RIPosome formation. **A:** Chemical structures of GSK583 and Gefitinib. **B:** Fluorescence micrographs of HeLa EGFP-RIPK2 cells treated for 30 min with 1 µM GSK583, 1 µM Gefitinib or DMSO as solvent control, following subsequent infection with *S. flexneri* M90T (right panel) or mock treatment (left panel). **C:** Immunoblot analysis of the cells in (B) probing for RIPK2 and RIPK2 pS176 to visualize RIPosome formation. **D:** IL-8 release in the supernatants from cells in (B and C) following infection with *S. flexneri* M90T after 2 h, 4 h and 6 h. Mean of triplicate measurements from one representative experiment with S.D. is shown.

## References

Abbott DW, Wilkins A, Asara JM, Cantley LC (2004) The Crohn’s disease protein, NOD2, requires RIP2 in order to induce ubiquitinylation of a novel site on NEMO. Curr Biol 14: 2217– 27

Andree M, Seeger JM, Schull S, Coutelle O, Wagner-Stippich D, Wiegmann K, Wunderlich CM, Brinkmann K, Broxtermann P, Witt A, Fritsch M, Martinelli P, Bielig H, Lamkemeyer T, Rugarli EI, Kaufmann T, Sterner-Kock A, Wunderlich FT, Villunger A, Martins LM et al. (2014) BID-dependent release of mitochondrial SMAC dampens XIAP-mediated immunity against Shigella. In EMBO J, pp 2171–87. England: 2014 The Authors.

Bertrand MJ, Doiron K, Labbe K, Korneluk RG, Barker PA, Saleh M (2009) Cellular inhibitors of apoptosis cIAP1 and cIAP2 are required for innate immunity signaling by the pattern recognition receptors NOD1 and NOD2. Immunity 30: 789–801

Bielig H, Lautz K, Braun PR, Menning M, Machuy N, Brugmann C, Barisic S, Eisler SA, Andree M, Zurek B, Kashkar H, Sansonetti PJ, Hausser A, Meyer TF, Kufer TA (2014) The cofilin phosphatase slingshot homolog 1 (SSH1) links NOD1 signaling to actin remodeling. In PLoS pathogens, p e1004351. United States:

Boyle JP, Parkhouse R, Monie TP (2014) Insights into the molecular basis of the NOD2 signalling pathway. Open biology 4

Browman DT, Resek ME, Zajchowski LD, Robbins SM (2006) Erlin-1 and erlin-2 are novel members of the prohibitin family of proteins that define lipid-raft-like domains of the ER. J Cell Sci 119: 3149–60

Carneiro LA, Travassos LH, Soares F, Tattoli I, Magalhaes JG, Bozza MT, Plotkowski MC, Sansonetti PJ, Molkentin JD, Philpott DJ, Girardin SE (2009) Shigella induces mitochondrial dysfunction and cell death in nonmyleoid cells. Cell Host Microbe 5: 123–36

Chamaillard M, Hashimoto M, Horie Y, Masumoto J, Qiu S, Saab L, Ogura Y, Kawasaki A, Fukase K, Kusumoto S, Valvano MA, Foster SJ, Mak TW, Nunez G, Inohara N (2003) An essential role for NOD1 in host recognition of bacterial peptidoglycan containing diaminopimelic acid. Nat Immunol 4: 702–7

Clerc P, Sansonetti PJ (1987) Entry of Shigella flexneri into HeLa cells: evidence for directed phagocytosis involving actin polymerization and myosin accumulation. Infect Immun 55: 2681–8

Damgaard RB, Fiil BK, Speckmann C, Yabal M, zur Stadt U, Bekker-Jensen S, Jost PJ, Ehl S, Mailand N, Gyrd-Hansen M (2013) Disease-causing mutations in the XIAP BIR2 domain impair NOD2-dependent immune signalling. EMBO molecular medicine 5: 1278–95

Damgaard RB, Nachbur U, Yabal M, Wong WW, Fiil BK, Kastirr M, Rieser E, Rickard JA, Bankovacki A, Peschel C, Ruland J, Bekker-Jensen S, Mailand N, Kaufmann T, Strasser A, Walczak H, Silke J, Jost PJ, Gyrd-Hansen M (2012) The ubiquitin ligase XIAP recruits LUBAC for NOD2 signaling in inflammation and innate immunity. Mol Cell 46: 746–58

Donnenberg MS, Donohue-Rolfe A, Keusch GT (1989) Epithelial cell invasion: an overlooked property of enteropathogenic Escherichia coli (EPEC) associated with the EPEC adherence factor. The Journal of infectious diseases 160: 452–459

Dorsch M, Wang A, Cheng H, Lu C, Bielecki A, Charron K, Clauser K, Ren H, Polakiewicz RD, Parsons T, Li P, Ocain T, Xu Y (2006) Identification of a regulatory autophosphorylation site in the serine-threonine kinase RIP2. Cell Signal 18: 2223–9

Dostert C, Meylan E, Tschopp J (2008) Intracellular pattern-recognition receptors. Adv Drug Deliv Rev 60: 830–40

Du C, Fang M, Li Y, Li L, Wang X (2000) Smac, a mitochondrial protein that promotes cytochrome c-dependent caspase activation by eliminating IAP inhibition. Cell 102: 33–42

Dupont A, Sommer F, Zhang K, Repnik U, Basic M, Bleich A, Kuhnel M, Backhed F, Litvak Y, Fulde M, Rosenshine I, Hornef MW (2016) Age-Dependent Susceptibility to Enteropathogenic Escherichia coli (EPEC) Infection in Mice. PLoS pathogens 12: e1005616

Feoktistova M, Geserick P, Kellert B, Dimitrova DP, Langlais C, Hupe M, Cain K, MacFarlane M, Hacker G, Leverkus M (2011) cIAPs block Ripoptosome formation, a RIP1/caspase-8 containing intracellular cell death complex differentially regulated by cFLIP isoforms. Molecular cell 43: 449–63

Fettweis G, Di Valentin E, L’Homme L, Lassence C, Dequiedt F, Fillet M, Coupienne I, Piette J (2017) RIP3 antagonizes a TSC2-mediated pro-survival pathway in glioblastoma cell death. Biochimica et biophysica acta 1864: 113–124

Gao W, Yang J, Liu W, Wang Y, Shao F (2016) Site-specific phosphorylation and microtubule dynamics control Pyrin inflammasome activation. Proceedings of the National Academy of Sciences of the United States of America 113: E4857–66

Garcia-Weber D, Dangeard AS, Cornil J, Thai L, Rytter H, Zamyatina A, Mulard LA, Arrieumerlou C (2018) ADP-heptose is a newly identified pathogen-associated molecular pattern of Shigella flexneri. EMBO reports

Gaudet RG, Guo CX, Molinaro R, Kottwitz H, Rohde JR, Dangeard AS, Arrieumerlou C, Girardin SE, Gray-Owen SD (2017) Innate Recognition of Intracellular Bacterial Growth Is Driven by the TIFA-Dependent Cytosolic Surveillance Pathway. Cell Rep 19: 1418–1430

Girardin SE, Boneca IG, Carneiro LA, Antignac A, Jehanno M, Viala J, Tedin K, Taha MK, Labigne A, Zahringer U, Coyle AJ, DiStefano PS, Bertin J, Sansonetti PJ, Philpott DJ (2003a) Nod1 detects a unique muropeptide from gram-negative bacterial peptidoglycan. Science 300: 1584–7

Girardin SE, Boneca IG, Viala J, Chamaillard M, Labigne A, Thomas G, Philpott DJ, Sansonetti PJ (2003b) Nod2 is a general sensor of peptidoglycan through muramyl dipeptide (MDP) detection. The Journal of biological chemistry 278: 8869–72

Girardin SE, Tournebize R, Mavris M, Page AL, Li X, Stark GR, Bertin J, DiStefano PS, Yaniv M, Sansonetti PJ, Philpott DJ (2001) CARD4/Nod1 mediates NF-kappaB and JNK activation by invasive Shigella flexneri. EMBO reports 2: 736–42

Goncharov T, Hedayati S, Mulvihill MM, Izrael-Tomasevic A, Zobel K, Jeet S, Fedorova AV, Eidenschenk C, deVoss J, Yu K, Shaw AS, Kirkpatrick DS, Fairbrother WJ, Deshayes K, Vucic D (2018) Disruption of XIAP-RIP2 Association Blocks NOD2-Mediated Inflammatory Signaling. Molecular cell 69: 551–565 e7

Gong Q, Long Z, Zhong FL, Teo DET, Jin Y, Yin Z, Boo ZZ, Zhang Y, Zhang J, Yang R, Bhushan S, Reversade B, Li Z, Wu B (2018) Structural basis of RIP2 activation and signaling. Nature communications 9: 4993

Haile PA, Votta BJ, Marquis RW, Bury MJ, Mehlmann JF, Singhaus R, Jr., Charnley AK, Lakdawala AS, Convery MA, Lipshutz DB, Desai BM, Swift B, Capriotti CA, Berger SB, Mahajan MK, Reilly MA, Rivera EJ, Sun HH, Nagilla R, Beal AM et al. (2016) The Identification and Pharmacological Characterization of 6-(tert-Butylsulfonyl)-N-(5-fluoro-1H-indazol-3-yl)quinolin-4-amine (GSK583), a Highly Potent and Selective Inhibitor of RIP2 Kinase. Journal of medicinal chemistry 59: 4867–80

Hall HT, Wilhelm MT, Saibil SD, Mak TW, Flavell RA, Ohashi PS (2008) RIP2 contributes to Nod signaling but is not essential for T cell proliferation, T helper differentiation or TLR responses. European journal of immunology 38: 64–72

Hara H, Tsuchiya K, Kawamura I, Fang R, Hernandez-Cuellar E, Shen Y, Mizuguchi J, Schweighoffer E, Tybulewicz V, Mitsuyama M (2013) Phosphorylation of the adaptor ASC acts as a molecular switch that controls the formation of speck-like aggregates and inflammasome activity. Nature immunology 14: 1247–55

Hasegawa M, Fujimoto Y, Lucas PC, Nakano H, Fukase K, Nunez G, Inohara N (2008) A critical role of RICK/RIP2 polyubiquitination in Nod-induced NF-kappaB activation. EMBO J 27: 373–83

Hausser A, Link G, Hoene M, Russo C, Selchow O, Pfizenmaier K (2006) Phospho-specific binding of 14-3-3 proteins to phosphatidylinositol 4-kinase III beta protects from dephosphorylation and stabilizes lipid kinase activity. J Cell Sci 119: 3613–21

Homer CR, Kabi A, Marina-Garcia N, Sreekumar A, Nesvizhskii AI, Nickerson KP, Chinnaiyan AM, Nunez G, McDonald C (2012) A dual role for receptor-interacting protein kinase 2 (RIP2) kinase activity in nucleotide-binding oligomerization domain 2 (NOD2)-dependent autophagy. The Journal of biological chemistry 287: 25565–76

Huang CC, Weng JH, Wei TY, Wu PY, Hsu PH, Chen YH, Wang SC, Qin D, Hung CC, Chen ST, Wang AH, Shyy JY, Tsai MD (2012) Intermolecular binding between TIFA-FHA and TIFA-pT mediates tumor necrosis factor alpha stimulation and NF-kappaB activation. Mol Cell Biol 32: 2664–73

Humphries F, Yang S, Wang B, Moynagh PN (2015) RIP kinases: key decision makers in cell death and innate immunity. Cell Death Differ 22: 225–36

Inohara N, Koseki T, Lin J, del Peso L, Lucas PC, Chen FF, Ogura Y, Nunez G (2000) An induced proximity model for NF-kappa B activation in the Nod1/RICK and RIP signaling pathways. The Journal of biological chemistry 275: 27823–31

Inohara N, Ogura Y, Fontalba A, Gutierrez O, Pons F, Crespo J, Fukase K, Inamura S, Kusumoto S, Hashimoto M, Foster SJ, Moran AP, Fernandez-Luna JL, Nunez G (2003) Host recognition of bacterial muramyl dipeptide mediated through NOD2. Implications for Crohn’s disease. J Biol Chem 278: 5509–12

Irving AT, Mimuro H, Kufer TA, Lo C, Wheeler R, Turner LJ, Thomas BJ, Malosse C, Gantier MP, Casillas LN, Votta BJ, Bertin J, Boneca IG, Sasakawa C, Philpott DJ, Ferrero RL, Kaparakis-Liaskos M (2014) The immune receptor NOD1 and kinase RIP2 interact with bacterial peptidoglycan on early endosomes to promote autophagy and inflammatory signaling. Cell Host Microbe 15: 623–35

Janeway CA, Jr. (1989) Approaching the asymptote? Evolution and revolution in immunology. Cold Spring Harb Symp Quant Biol 54 Pt 1: 1–13

Jeru I, Papin S, L’Hoste S, Duquesnoy P, Cazeneuve C, Camonis J, Amselem S (2005) Interaction of pyrin with 14.3.3 in an isoform-specific and phosphorylation-dependent manner regulates its translocation to the nucleus. Arthritis Rheum 52: 1848–57

Jo Y, Sguigna PV, DeBose-Boyd RA (2011) Membrane-associated ubiquitin ligase complex containing gp78 mediates sterol-accelerated degradation of 3-hydroxy-3-methylglutaryl-coenzyme A reductase. The Journal of biological chemistry 286: 15022–31

Kashkar H, Seeger JM, Hombach A, Deggerich A, Yazdanpanah B, Utermohlen O, Heimlich G, Abken H, Kronke M (2006) XIAP targeting sensitizes Hodgkin lymphoma cells for cytolytic T-cell attack. Blood 108: 3434–40

Keestra-Gounder AM, Byndloss MX, Seyffert N, Young BM, Chavez-Arroyo A, Tsai AY, Cevallos SA, Winter MG, Pham OH, Tiffany CR, de Jong MF, Kerrinnes T, Ravindran R, Luciw PA, McSorley SJ, Baumler AJ, Tsolis RM (2016) NOD1 and NOD2 signalling links ER stress with inflammation. Nature

Keller A, Nesvizhskii AI, Kolker E, Aebersold R (2002) Empirical statistical model to estimate the accuracy of peptide identifications made by MS/MS and database search. Anal Chem 74: 5383–92

Krieg A, Correa RG, Garrison JB, Le Negrate G, Welsh K, Huang Z, Knoefel WT, Reed JC (2009) XIAP mediates NOD signaling via interaction with RIP2. Proceedings of the National Academy of Sciences of the United States of America 106: 14524–9

Kufer TA (2008) Signal transduction pathways used by NLR-type innate immune receptors. Molecular bioSystems 4: 380–6

Kufer TA, Kremmer E, Adam AC, Philpott DJ, Sansonetti PJ (2008) The pattern-recognition molecule Nod1 is localized at the plasma membrane at sites of bacterial interaction. Cellular microbiology 10: 477–486

Kufer TA, Kremmer E, Banks DJ, Philpott DJ (2006) Role for erbin in bacterial activation of Nod2. Infect Immun 74: 3115–24

Lauer SA, Iyer S, Sanchez T, Forst CV, Bowden B, Carlson K, Sriranganathan N, Boyle SM (2014) Proteomic analysis of detergent resistant membrane domains during early interaction of macrophages with rough and smooth Brucella melitensis. PLoS one 9: e91706

Lecine P, Esmiol S, Metais JY, Nicoletti C, Nourry C, McDonald C, Nunez G, Hugot JP, Borg JP, Ollendorff V (2007) The NOD2-RICK complex signals from the plasma membrane. In The Journal of biological chemistry, pp 15197–207. United States:

Lembo-Fazio L, Nigro G, Noel G, Rossi G, Chiara F, Tsilingiri K, Rescigno M, Rasola A, Bernardini ML (2011) Gadd45alpha activity is the principal effector of Shigella mitochondria-dependent epithelial cell death in vitro and ex vivo. Cell Death Dis 2: e122

Liu HM, Loo YM, Horner SM, Zornetzer GA, Katze MG, Gale M, Jr. (2012) The mitochondrial targeting chaperone 14-3-3epsilon regulates a RIG-I translocon that mediates membrane association and innate antiviral immunity. Cell host & microbe 11: 528–37

Maharana J, Pradhan SK, De S (2017) NOD1CARD Might Be Using Multiple Interfaces for RIP2-Mediated CARD-CARD Interaction: Insights from Molecular Dynamics Simulation. PLoS one 12: e0170232

Manon F, Favier A, Nunez G, Simorre JP, Cusack S (2007) Solution structure of NOD1 CARD and mutational analysis of its interaction with the CARD of downstream kinase RICK. J Mol Biol 365: 160–74

Mayle S, Boyle JP, Sekine E, Zurek B, Kufer TA, Monie TP (2014) Engagement of nucleotide-binding oligomerization domain-containing protein 1 (NOD1) by receptor-interacting protein 2 (RIP2) is insufficient for signal transduction. The Journal of biological chemistry 289: 22900–14

McCarthy JV, Ni J, Dixit VM (1998) RIP2 is a novel NF-kappaB-activating and cell death-inducing kinase. The Journal of biological chemistry 273: 16968–75

Moghaddas F, Llamas R, De Nardo D, Martinez-Banaclocha H, Martinez-Garcia JJ, Mesa-Del-Castillo P, Baker PJ, Gargallo V, Mensa-Vilaro A, Canna S, Wicks IP, Pelegrin P, Arostegui JI, Masters SL (2017) A novel Pyrin-Associated Autoinflammation with Neutrophilic Dermatosis mutation further defines 14-3-3 binding of pyrin and distinction to Familial Mediterranean Fever. Ann Rheum Dis 76: 2085–2094

Nachbur U, Stafford CA, Bankovacki A, Zhan Y, Lindqvist LM, Fiil BK, Khakham Y, Ko HJ, Sandow JJ, Falk H, Holien JK, Chau D, Hildebrand J, Vince JE, Sharp PP, Webb AI, Jackman KA, Muhlen S, Kennedy CL, Lowes KN et al. (2015) A RIPK2 inhibitor delays NOD signalling events yet prevents inflammatory cytokine production. Nature communications 6: 6442

Nakamura N, Lill JR, Phung Q, Jiang Z, Bakalarski C, de Maziere A, Klumperman J, Schlatter M, Delamarre L, Mellman I (2014) Endosomes are specialized platforms for bacterial sensing and NOD2 signalling. Nature 509: 240–4

Navas TA, Baldwin DT, Stewart TA (1999) RIP2 is a Raf1-activated mitogen-activated protein kinase kinase. The Journal of biological chemistry 274: 33684–90

Ogura Y, Inohara N, Benito A, Chen FF, Yamaoka S, Nunez G (2001) Nod2, a Nod1/Apaf-1 family member that is restricted to monocytes and activates NF-kappaB. J Biol Chem 276: 4812–8

Panda S, Gekara NO (2018) The deubiquitinase MYSM1 dampens NOD2-mediated inflammation and tissue damage by inactivating the RIP2 complex. Nature communications 9: 4654

Pearce MM, Wang Y, Kelley GG, Wojcikiewicz RJ (2007) SPFH2 mediates the endoplasmic reticulum-associated degradation of inositol 1,4,5-trisphosphate receptors and other substrates in mammalian cells. The Journal of biological chemistry 282: 20104–15

Pearce MM, Wormer DB, Wilkens S, Wojcikiewicz RJ (2009) An endoplasmic reticulum (ER) membrane complex composed of SPFH1 and SPFH2 mediates the ER-associated degradation of inositol 1,4,5-trisphosphate receptors. The Journal of biological chemistry 284: 10433–45

Pellegrini E, Desfosses A, Wallmann A, Schulze WM, Rehbein K, Mas P, Signor L, Gaudon S, Zenkeviciute G, Hons M, Malet H, Gutsche I, Sachse C, Schoehn G, Oschkinat H, Cusack S (2018) RIP2 filament formation is required for NOD2 dependent NF-kappaB signalling. Nature communications 9: 4043

Pellegrini E, Signor L, Singh S, Boeri Erba E, Cusack S (2017) Structures of the inactive and active states of RIP2 kinase inform on the mechanism of activation. PLoS one 12: e0177161

Pozuelo Rubio M, Geraghty KM, Wong BH, Wood NT, Campbell DG, Morrice N, Mackintosh C (2004) 14-3-3-affinity purification of over 200 human phosphoproteins reveals new links to regulation of cellular metabolism, proliferation and trafficking. Biochem J 379: 395–408

Riedl J, Crevenna AH, Kessenbrock K, Yu JH, Neukirchen D, Bista M, Bradke F, Jenne D, Holak TA, Werb Z, Sixt M, Wedlich-Soldner R (2008) Lifeact: a versatile marker to visualize F-actin. Nat Methods 5: 605–7

Ruefli-Brasse AA, Lee WP, Hurst S, Dixit VM (2004) Rip2 participates in Bcl10 signaling and T-cell receptor-mediated NF-kappaB activation. The Journal of biological chemistry 279: 1570–4

Sprenger RR, Fontijn RD, van Marle J, Pannekoek H, Horrevoets AJ (2006) Spatial segregation of transport and signalling functions between human endothelial caveolae and lipid raft proteomes. Biochem J 400: 401–10

Sprenger RR, Speijer D, Back JW, De Koster CG, Pannekoek H, Horrevoets AJ (2004) Comparative proteomics of human endothelial cell caveolae and rafts using two-dimensional gel electrophoresis and mass spectrometry. Electrophoresis 25: 156–72

Tao M, Scacheri PC, Marinis JM, Harhaj EW, Matesic LE, Abbott DW (2009) ITCH K63-ubiquitinates the NOD2 binding protein, RIP2, to influence inflammatory signaling pathways. Curr Biol 19: 1255–63

Tattoli I, Sorbara MT, Vuckovic D, Ling A, Soares F, Carneiro LA, Yang C, Emili A, Philpott DJ, Girardin SE (2012) Amino acid starvation induced by invasive bacterial pathogens triggers an innate host defense program. Cell host & microbe 11: 563–75

Tenev T, Bianchi K, Darding M, Broemer M, Langlais C, Wallberg F, Zachariou A, Lopez J, MacFarlane M, Cain K, Meier P (2011) The Ripoptosome, a signaling platform that assembles in response to genotoxic stress and loss of IAPs. Molecular cell 43: 432–48

Thome M, Hofmann K, Burns K, Martinon F, Bodmer JL, Mattmann C, Tschopp J (1998) Identification of CARDIAK, a RIP-like kinase that associates with caspase-1. Curr Biol 8: 885–8

Tigno-Aranjuez JT, Asara JM, Abbott DW (2010) Inhibition of RIP2’s tyrosine kinase activity limits NOD2-driven cytokine responses. Genes Dev 24: 2666–77

Tigno-Aranjuez JT, Benderitter P, Rombouts F, Deroose F, Bai X, Mattioli B, Cominelli F, Pizarro TT, Hoflack J, Abbott DW (2014) In vivo inhibition of RIPK2 kinase alleviates inflammatory disease. The Journal of biological chemistry 289: 29651–64

Ting JP, Lovering RC, Alnemri ES, Bertin J, Boss JM, Davis BK, Flavell RA, Girardin SE, Godzik A, Harton JA, Hoffman HM, Hugot JP, Inohara N, Mackenzie A, Maltais LJ, Nunez G, Ogura Y, Otten LA, Philpott D, Reed JC et al. (2008) The NLR gene family: a standard nomenclature. Immunity 28: 285–7

Todd AG, Morse R, Shaw DJ, Stebbings H, Young PJ (2010) Analysis of SMN-neurite granules: Core Cajal body components are absent from SMN-cytoplasmic complexes. Biochem Biophys Res Commun 397: 479–85

Travassos LH, Carneiro LA, Ramjeet M, Hussey S, Kim YG, Magalhaes JG, Yuan L, Soares F, Chea E, Le Bourhis L, Boneca IG, Allaoui A, Jones NL, Nunez G, Girardin SE, Philpott DJ (2010) Nod1 and Nod2 direct autophagy by recruiting ATG16L1 to the plasma membrane at the site of bacterial entry. Nature immunology 11: 55–62

UniProt Consortium T (2018) UniProt: the universal protein knowledgebase. Nucleic Acids Res 46: 2699

Wang Y, Pearce MM, Sliter DA, Olzmann JA, Christianson JC, Kopito RR, Boeckmann S, Gagen C, Leichner GS, Roitelman J, Wojcikiewicz RJ (2009) SPFH1 and SPFH2 mediate the ubiquitination and degradation of inositol 1,4,5-trisphosphate receptors in muscarinic receptor-expressing HeLa cells. Biochimica et biophysica acta 1793: 1710–8

Warner N, Burberry A, Franchi L, Kim YG, McDonald C, Sartor MA, Nunez G (2013) A genome-wide siRNA screen reveals positive and negative regulators of the NOD2 and NF-kappaB signaling pathways. Science signaling 6: rs3

Warner N, Burberry A, Pliakas M, McDonald C, Nunez G (2014) A genome-wide small interfering RNA (siRNA) screen reveals nuclear factor-kappaB (NF-kappaB)-independent regulators of NOD2-induced interleukin-8 (IL-8) secretion. The Journal of biological chemistry 289: 28213–24

Windheim M, Lang C, Peggie M, Plater LA, Cohen P (2007) Molecular mechanisms involved in the regulation of cytokine production by muramyl dipeptide. Biochem J 404: 179–90

Witt A, Vucic D (2017) Diverse ubiquitin linkages regulate RIP kinases-mediated inflammatory and cell death signaling. Cell Death Differ 24: 1160–1171

Yabal M, Muller N, Adler H, Knies N, Gross CJ, Damgaard RB, Kanegane H, Ringelhan M, Kaufmann T, Heikenwalder M, Strasser A, Gross O, Ruland J, Peschel C, Gyrd-Hansen M, Jost PJ (2014) XIAP restricts TNF-and RIP3-dependent cell death and inflammasome activation. Cell Rep 7: 1796–808

Zhang DW, Shao J, Lin J, Zhang N, Lu BJ, Lin SC, Dong MQ, Han J (2009) RIP3, an energy metabolism regulator that switches TNF-induced cell death from apoptosis to necrosis. Science 325: 332–6

Zurek B, Proell M, Wagner RN, Schwarzenbacher R, Kufer TA (2012) Mutational analysis of human NOD1 and NOD2 NACHT domains reveals different modes of activation. Innate Immun 18: 100–11

